# Coexpression of CCR7 and CXCR4 during B cell development controls CXCR4 responsiveness and bone marrow homing

**DOI:** 10.1101/689372

**Authors:** Saria Mcheik, Nils Van Eeckhout, Cédric De Poorter, Céline Galés, Marc Parmentier, Jean-Yves Springael

## Abstract

G protein-coupled receptors (GPCR) constitute the largest family of plasma membrane proteins involved in cell signaling. Besides their canonical role in signaling, GPCR can also act as allosteric modulator of one another through receptor oligomerization. However, only few studies have investigated the relevance of GPCR oligomerization, and the role of GPCR interaction in physiological processes remains largely unknown. By using chemokine receptors and B cell development as a model system, we unveil in this study a novel role for CCR7 as a selective endogenous allosteric modulator of CXCR4. We show that the upregulation of CCR7 expression naturally occurring in late stages of B cell development contributes to the functional inactivation of CXCR4, and that B cell populations from CCR7^-/-^ mice display higher responsiveness to CXCL12 and improved retention in the bone marrow parenchyma. We also provide molecular evidences supporting a model in which upregulation of CCR7 favors the formation of CXCR4-CCR7 heteromers wherein CXCR4 is selectively impaired in its ability to activate some G protein complexes.

## Introduction

G protein-coupled receptors (GPCRs) are a large family of receptor proteins that respond to a great range of extracellular stimuli. They are involved in many physiological processes and are targeted by approximately 40% of all medicinal drugs [1]. Over the last decade, it has become widely accepted that most GPCRs possess, besides their orthosteric site, spatially distinct allosteric sites that can modulate the receptors function [2, 3]. There is also evidences that GPCRs have the potential to act as endogenous allosteric modulators of one another through receptor oligomerization. Oligomerization is reported to influence many aspects of receptor function, including modulation of the ligand occupancy, change in the selectivity of one receptor for a particular ligand, modification of the signal efficacy or complete signal switching [4, 5]. Nevertheless, the physiological relevance of GPCRs oligomerization remains yet to be determined. By using chemokines receptors as a model system, we previously shown that CXCR4 heteromerization with CCR2 or CCR5 results in negative binding cooperativity of allosteric nature in leukocyte populations that coexpress these receptors endogenously [6, 7]. With the aim of studying the consequences of CXCR4 heteromerization in a physiological process, we investigate the putative allostery between CXCR4 and CCR7 during B cells development. CXCR4 is expressed at all stages of B cell development in bone marrow from HSCs to mature B cells and plays a major role in the homing of B cell precursors [8,9,10,11]. When lymphopoiesis progress to more differentiated stages, B cells lose their responsiveness to CXCL12 despite the continuous expression of CXCR4 [12,13,14,15,9]. B cells regain their sensitivity to CXCL12 as they further differentiate into plasma cells, but the mechanism involved in this transient loss of CXCR4 responsiveness was not identified. B cells differentiation in bone marrow is accompanied by the modulation of expression of several receptors. Amongst them, CCR7 is strongly upregulated during the transition of Pre-B cells to immature and mature stages [16]. CCR7 is required, together with CXCR4 and CXCR5, for B cells entry to lymphoid organs that express high levels of chemokines CCL19 and CCL21. As CCR7 upregulation takes place in B cell populations known to display poor CXCR4 responsiveness, we investigated whether CCR7 could be involved in the inhibition of CXCR4 function.

In the present study, we identified a novel regulatory function associated to CCR7 expression, showing that CCR7 controls the responsiveness of CXCR4 and the homing of B cells into bone marrow. Using a combination of approaches, we demonstrate that CCR7 physically interacts with CXCR4 and that CCR7/CXCR4 heteromers exhibit reduced CXCR4 signaling capacity compared to CXCR4 homomers.

## Results

### CCR7 inhibits CXCR4 responsiveness during B cell development

In order to evaluate the influence of CCR7 on CXCR4 function, we first tested the expression and functional response of the two receptors in various B cell populations from WT and CCR7^-/-^ mice. Bone marrow cells were sorted into three subpopulations according to established markers and the expression of chemokine receptors was measured by RT- qPCR [17, 18] (Fig 1A and S1A Fig). In agreement with previous studies, we showed that CXCR4 was expressed in pre-B cells and that its expression decreased about three-fold in immature and mature B cells. In contrast, expression of CCR7 was weak in pre-B cells and increased about two-fold as differentiation progressed to immature and mature B cells. Finally, expression of CXCR5 and CCR6 was barely detectable in pre-B cells and increased in immature and mature B cells. By using FACS, we confirmed that CXCR4 was expressed at the surface of pre-B cells and that its expression decreased in immature and mature B cells (Fig 1B). By comparison, cell surface expression of CCR7 was weak in pre-B cells but increased as differentiation progressed to immature and mature stages. In agreement with the RT-qPCR experiments, the expression of CXCR5 and CCR6 at the cell surface was only detectable in immature and mature B cells (Fig 1B). As CCR7 upregulation at the cell surface takes place in populations known to display poor responsiveness to CXCR4 agonists, we wondered whether it might be involved in the impairment of CXCR4 activity. We first investigated the impact of CCR7 expression on the presence of CXCR4 at the cell surface and showed that the signal for CXCR4, as well as for CCR6 and CXCR5, was similar for populations from CCR7^-/-^ or control mice (Fig 1B). Next, we investigated the impact of CCR7-deficiency on the responsiveness of CXCR4 by measuring the ability of B cells to migrate *ex vivo* towards a CXCL12 gradient. In agreement with previous studies [12,13,14,15,9], chemotaxis of B cells from CCR7^+/+^ wild-type (WT) mice decreased as differentiation progressed, the mature B cells being almost unresponsive to CXCL12 (Fig 1C, 1D and S2 Fig). In contrast, mature B cells from CCR7^-/-^ mice migrated significantly more efficiently than control cells (Fig 1C, 1D S1 and S2 Fig). A higher migration index of immature B cells from CCR7^-/-^ mice was also observed although the difference did not reach statistical significance. An improved CXCR4 responsiveness was also detected in CCR7^-/-^ mature B cells by using receptor internalization or calcium mobilization assays (S3 Fig). The migration of CCR7-deficient mature B cells was completely abrogated upon pre-treatment with the CXCR4-selective antagonist AMD3100 or the blocking monoclonal MAB21625, confirming the involvement of CXCR4 (Fig 1E). In contrast, CCR7 blockade with monoclonal antibody MAB3477 did not restore CXCR4 responsiveness to CCR7^+/+^ mature B cells, indicating that CCR7 signaling is not required (Fig 1F). Importantly, CCR7-deficiency did not increase the responsiveness of CXCR5 or CCR6, suggesting that CCR7 controls selectively the function of CXCR4 (Fig 1G).

**Figure 1.**
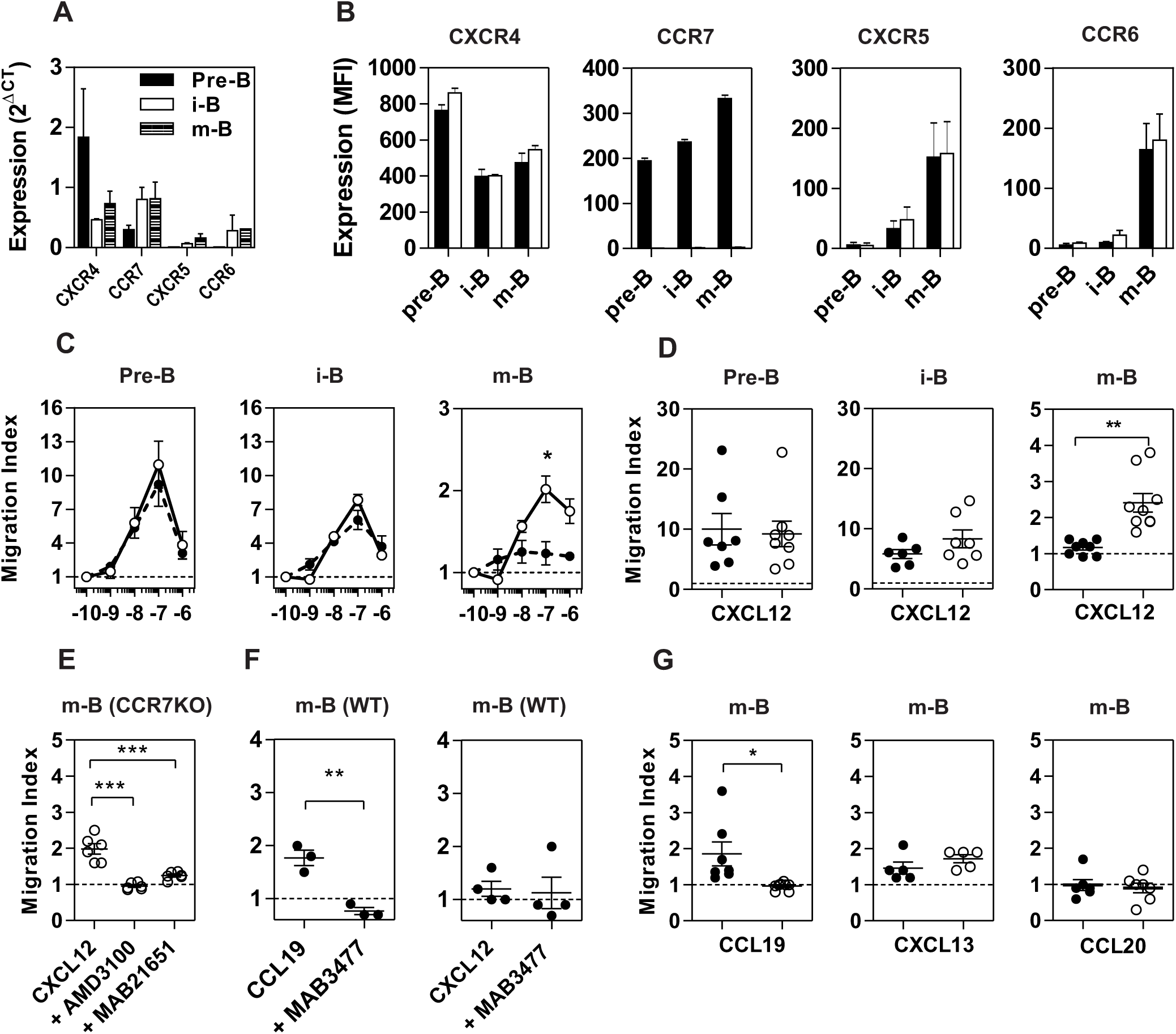
Properties of B cell populations prepared from CCR7^+/+^ or CCR7^-/-^ mice. **A. Expression of chemokine receptors in B cell subpopulations.** Bone marrow B cell subpopulations were discriminated and sorted by flow cytometry and the expression of CXCR4, CCR7, CXCR5 and CCR6 was quantified in each B cell subpopulation by RT-qPCR using GAPDH and β-actin as references. **B. Cell surface expression of chemokine receptors.** The cell surface expression of chemokine receptors in each B cell subpopulation was estimated by flow cytometry using PE-conjugated anti-CXCR4 and APC-conjugated anti-CCR7, anti-CXCR5 and anti-CCR6 antibodies. Bars represent mean values ± SEM (n = 5) of the mean fluorescence index for each receptor, detected in Pre-B, immature (i-B) or mature B cells (m-B) from CCR7^+/+^ (black bars) or CCR7^-/-^ mice (white bars). **C. Chemotaxis of bone marrow B cells toward CXCL12.** Transwell migration of bone marrow B cells from CCR7^+/+^ (black symbols) or CCR7^-/-^ (white symbols) mice in response to increasing concentrations of CXCL12. Migration index after a 2-h incubation were plotted for Pre-(B220^+^/IgM^-^/IgD^-^), immature (B220^+^/IgM^+^/IgD^-^) and mature (B220^+^/IgM^+^/IgD^+^) B cells. All conditions were run in triplicates and the data, representative of two independent experiments, are presented as mean values ± SEM, *, P < 0.05). **D. Enhanced chemotactic response to CXCL12 in mature B cells from CCR7^-/-^ mice.** Transwell migration of bone marrow B cells from CCR7^+/+^ (black dots) and CCR7^-/-^ (white dots) mice in response to 100 nM CXCL12. Migration indexes were plotted for each subpopulation. Data are represented as mean values ± SEM and dots correspond to individual mice (n = 6 to 8; **, P < 0.005). **E. Blockade of CXCR4 inhibits CXCL12-elicited migration of CCR7^-/-^ mature B cells.** Transwell migration of bone marrow mature B cells from CCR7^-/-^ mice in response to 100 nM CXCL12 in the presence or not of 1 µM AMD3100 or 5 µg/ml MAB2165 (blocking anti-CXCR4 monoclonal). Data are presented as mean values ± SEM and dots correspond to individual mice (n = 6; ***, P < 0.0005) **F. Blockade of CCR7 does not affect CXCR4 responsiveness in CCR7^+/+^ mature B cells.** Transwell migration of bone marrow mature B cells from CCR7^+/+^ mice in response to 100 nM CCL19 or CXCL12, and in the presence or not of 5 µg/ml MAB3477 (blocking anti-CCR7 monoclonal). Data are presented as mean values ± SEM, and dots correspond to individual mice (n = 3 to 4; **, P < 0.005) **G. CCR7-deficiency abrogates the CCL19-dependent migration of mature B cells but does not affect the migration in response to CXCL13 or CCL20.** Transwell migration of bone marrow mature B cells from CCR7^+/+^ (black dots) and CCR7^-/-^ (white dots) mice in response to 100 nM CCL19, CXCL13 or CCL20. Data are represented as mean values ± SEM, and dots correspond to individual mice (n = 5 to 8; *, P < 0.05).

### CCR7 regulates the number of mature B cells in bone marrow

Because CXCR4 plays a major role in the homing of progenitors to bone marrow, we measured the number of B cells in mice expressing or not CCR7, and found that the total number of B220^+^ B cells is increased moderately in the bone marrow of CCR7^-/-^ mice (S4 Fig). When analyzing the proportions of B cell subpopulations, we showed that the number of mature B cells is selectively increased in the bone marrow of CCR7^-/-^ mice (Fig 2A, and S5 Fig) while the number of CD4^+^ or CD8^+^ cells remains similar in CCR7^-/-^ and wild type mice (S6 Fig). We next performed an in vivo pulse labeling experiment to discriminate B cell subpopulations present in bone marrow parenchyma and sinusoids [19]. We injected mice with a phycoerythin-conjugated CD19 antibody shortly before sacrifice, and analyzed B cell subpopulations in sinusoids (CD19^+^) and parenchyma (CD19^-^). An increase in the proportion of immature and mature B cells was found in the parenchyma of CCR7^-/-^ mice, while the number of these cells decreased significantly in bone marrow sinusoids (Fig. 2B and 2C). Yet, the number of immature and mature B cells in blood was similar in CCR7^-/-^ and wild type mice, in agreement with previous studies (Fig 2D) [20]. Collectively, these data indicate that the regulation of CXCR4 responsiveness by CCR7 impacts selectively the retention of immature and mature B cells in the parenchymal compartment of bone marrow. To further analyze the contribution of CCR7 to B cell migration, we performed an adoptive transfer experiments. We compared the ability of B cells to migrate and home to bone marrow by fluorescently labeling WT and CCR7^-/-^ B220^+^ cells purified from bone marrow and transferring them into WT mice. The CCR7-deficiency increases about two-fold the homing of transferred immature B cells to bone marrow (Fig 3). The CXCR4-selective antagonist AMD3100 inhibited drastically the number of cells recovered in bone marrow, confirming the role of CXCR4 in the migration/retention process of immature B cells (Fig 3). In contrast, mature B cells populated bone marrow with similar efficiency whether the cells expressed or not CCR7, which may appear paradoxical in the light of the results generated in *ex vivo* chemotaxis assays. However, control experiments performed in the presence of AMD3100 revealed that CXCR4 does not play a major role in the homing of mature B cells *in vivo*, ruling out the relevance of investigating CCR7 regulatory function on mature B cells. Next, we used bone marrow cells from CCR7^-/-^ or WT CD45.2^+^ mice to reconstitute the hematopoietic compartment of irradiated WT CD45.1^+^ mice (Fig 4A). We showed that the number of CCR7^-/-^ B220^+^ cells is reduced in the bone marrow of reconstituted mice. However, when analyzing B cell subpopulations, we found that the number of mature B cells remains unaffected while the number of Pre B cells decreases significantly, which results in a higher proportion of CCR7^-/-^ mature B cells in the bone marrow of reconstituted mice. However, whether this observation is due to a more efficient retention of mature B cells or a direct effect of CCR7 on hematopoiesis remains to be determined. We also generated reverse chimeras where CCR7^-/-^ and WT CD45.2^+^ mice were irradiated and reconstituted with the bone marrow of WT CD45.1^+^ mice (Figure 4B). In this setting, the number of B cells remained similar in both groups, suggesting that increased retention of B cells in the bone marrow of CCR7^-/-^ mice is not due to an alteration of the bone marrow microenvironment.

**Figure 2.**
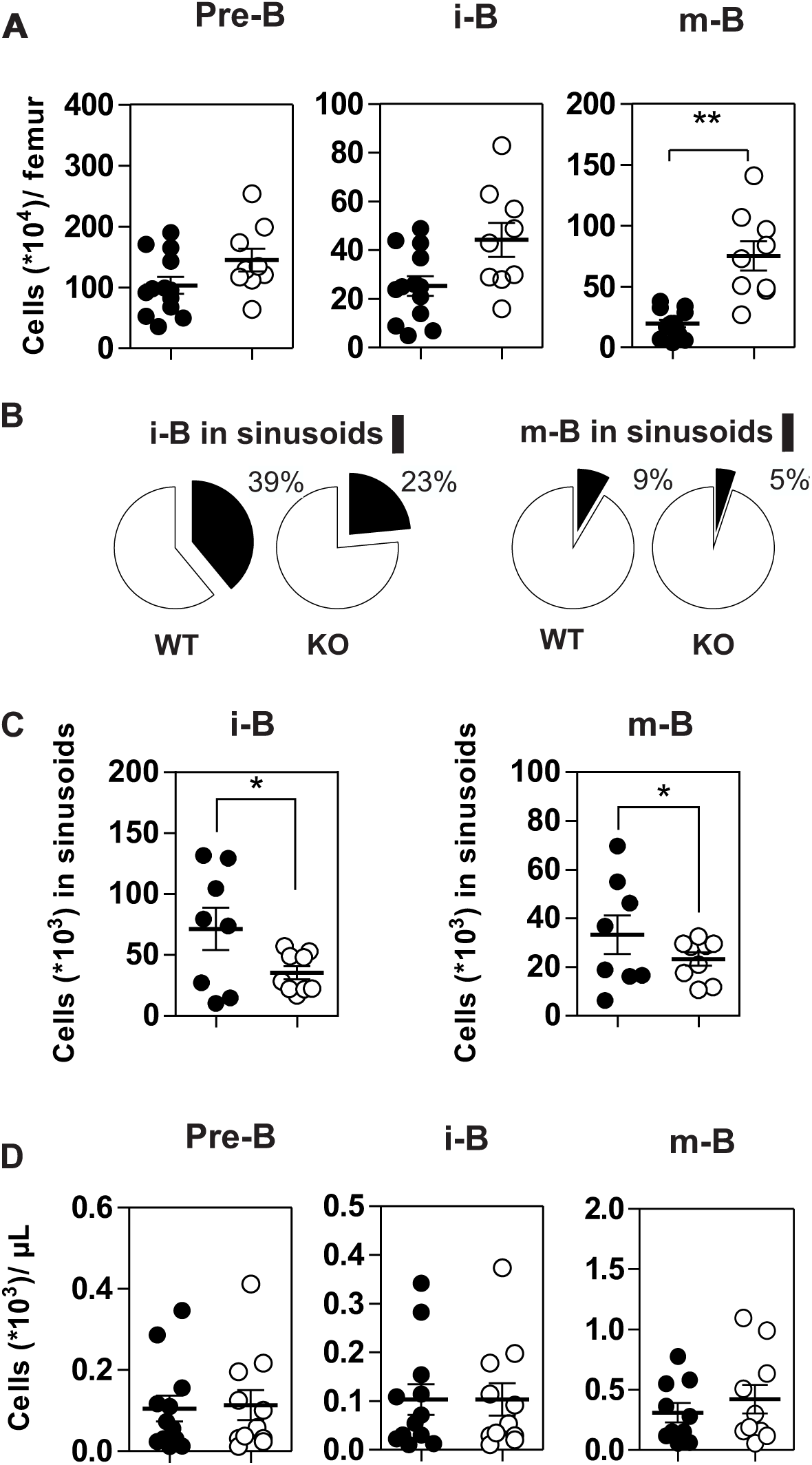
**A. Increased number of mature B cells in the bone marrow of CCR7^-/-^ mice.** The number of B cell subpopulations was determined in the bone marrow of CCR7^+/+^ (black dots) and CCR7^-/-^ (white dots) mice using AF700-conjugated anti-B220, FITC-conjugated anti-CD43, PerCP Cy5.5-conjugated anti-IgM and V450-conjugated anti-IgD antibodies. Data are represented as mean values ± SEM and dots correspond to individual mice (n = 10 to 13 mice; *, P<0.05; **, P < 0.005). **B-C Decreased egress of immature and mature B cells in the bone marrow sinusoids of CCR7^-/-^ mice**. In vivo labelling of bone marrow B cell subsets by injection of PE-conjugated anti-CD19 antibodies 4 minutes before sacrifice and tissue collection. The proportions of immature and mature B cells in the parenchyma (CD19^-^; white) and sinusoids (CD19+; black) are displayed in **B** and the number of immature and mature B cells in sinusoids displayed in **C**. Data for CCR7^+/+^ (black dots) and CCR7^-/-^ (white dots) mice are represented as mean values ± SEM and dots correspond to individual mice (n = 8 to 9 mice; *, P<0.05). **D. B cell counts are comparable in the blood of CCR7^+/+^ and CCR7^-/-^ mice.** The number of B cell subpopulations was determined in the blood of CCR7^+/+^ (black dots) and CCR7^-/-^ (white dots) mice using AF700-conjugated anti-B220, FITC-conjugated anti-CD43, PerCP Cy5.5-conjugated anti-IgM and V450-conjugated anti-IgD antibodies. Data are represented as mean values ± SEM and dots correspond to individual mice (n = 10 to 13 mice; *, P<0.05; **, P < 0.005).

**Figure 3.**
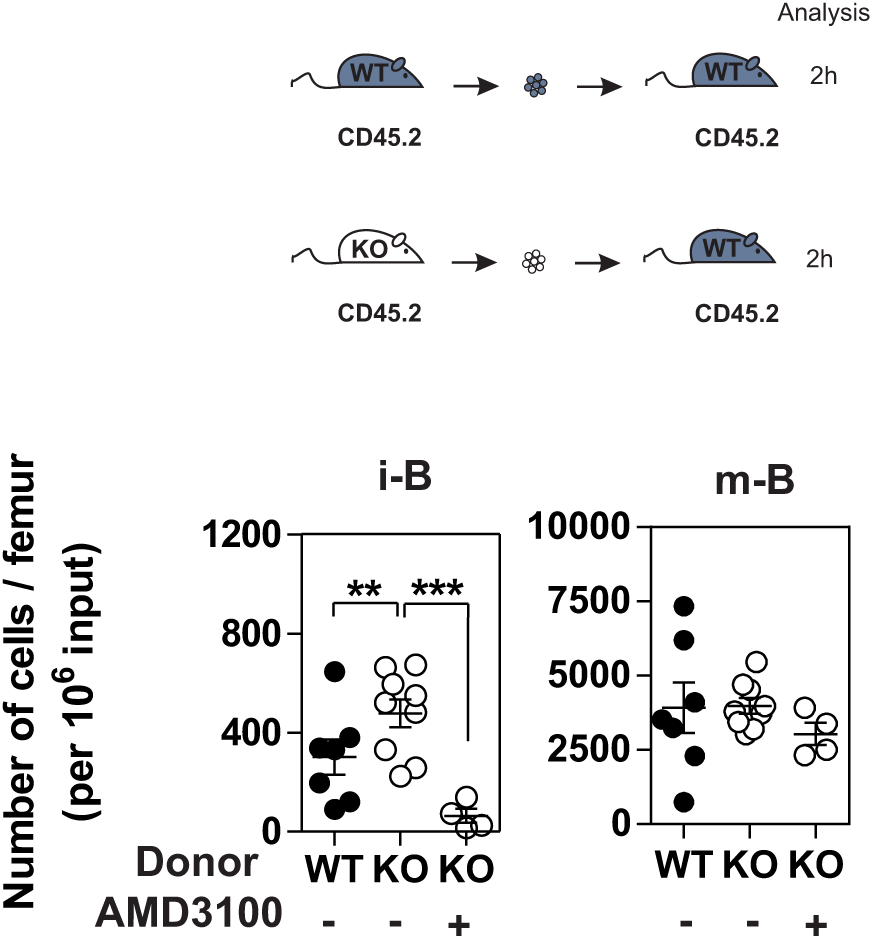
CCR7 deficiency results in increased homing of immature B cells to bone marrow. B220^+^ cells purified from bone marrow of CCR7^+/+^ (black dots) or CCR7^-/-^ mice (white dots) were labeled with CFDA-SE and transferred into wild type recipients by injection into the retro-orbital venous plexus in combination or not with AMD3100. Two hours later, bone marrow mononuclear cells were isolated and B cell subpopulations were analyzed by flow cytometry using AF700-conjugated anti-B220, PerCP Cy5.5-conjugated anti-IgM and V450-conjugated anti-IgD antibodies. Data are represented as mean values ± SEM and dots correspond to individual mice (n = 4 to 8 mice; **, P < 0.005; ***, P<0.0005).

**Figure 4.**
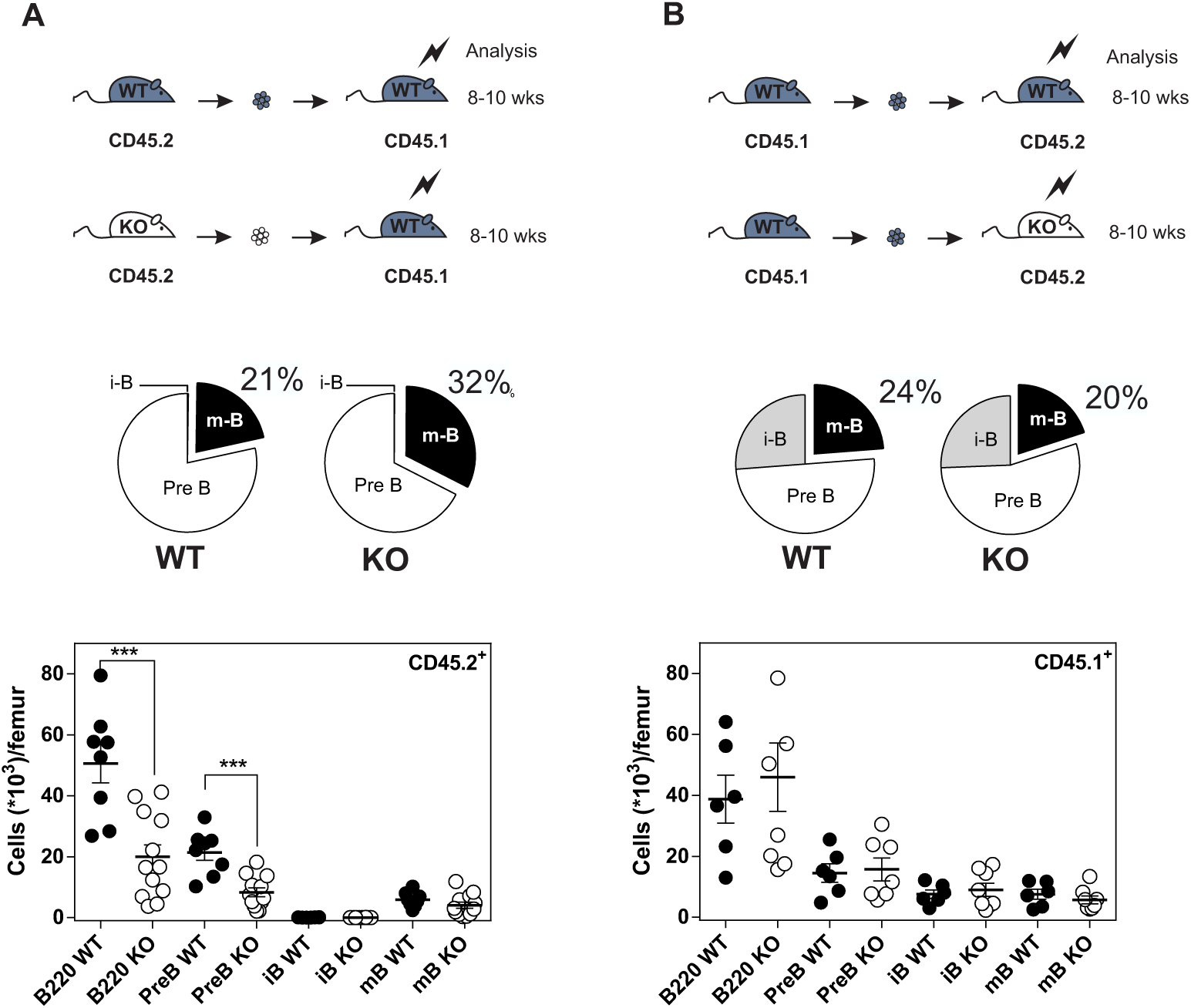
Increased number of B cells in the bone marrow of CCR7^-/-^ mice is not due to an alteration of bone marrow environment. **A.** Distribution and quantification of donor CD45.2^+^ (WT or CCR7^-/-^) B cells recovered from the bone marrow of chimeric CD45.1^+^ recipients. **B.** Distribution and quantification of donor CD45.1^+^ WT B cells recovered from the bone marrow of chimeric CD45.2^+^ (WT or CCR7^-/-^) recipients. Data are represented as mean values ± SEM and dots correspond to individual mice (n = 8 to 12 mice; P<0.0005).

### Regulation of CXCR4 responsiveness does not require CCR7 signaling

To get further insight into the mechanisms underlying the inhibition of CXCR4 responsiveness in B cells, we overexpressed CCR7 in the human pre-B cell line Nalm-6. While expression of CCR7 did not affect the presence of CXCR4 at the cell surface (Fig 4A), it induced a strong inhibition of CXCR4 responsiveness as measured in cell adhesion (Fig 4B) or chemotaxis (Fig 4C and 4D) assays. By comparison, expression of CCR5 did not impact on CXCR4 function (Fig 4D). Thus, expression of CCR7 in a human pre-B cell line seems to be sufficient to mimic the inhibition of CXCR4 responsiveness detected during normal B cell development *in vivo*. Then, we took advantage of this expression system to investigate the effect of CCR7 mutants unable to bind chemokines (CCR7^ΔNT^) or to activate signaling pathways (CCR7^(N/A)PXXY^ and CCR7^D(R/A)Y^). Chemotaxis experiments showed that the three non-functional CCR7 mutants inhibit CXCR4 responsiveness as efficiently as wild type CCR7 (Fig 4D), confirming that signaling of CCR7 is not required for its inhibitory activity on CXCR4 function.

### CCR7 inhibits CXCR4-mediated Gαi proteins activation

To better understand the regulation of CXCR4 responsiveness by CCR7, we measured the ability of CXCR4 to activate G proteins using Bioluminescence Resonance Energy Transfer (BRET). This technology was previously shown to provide an accurate sensitivity to probe the activation of specific G protein isoforms [21,22,23]. The BRET-based G protein activation assay measures the conformational change occurring within G proteins upon GDP/GTP exchange, which is sensed as a decrease in the BRET^2^ signal between the probes fused to Gαi and Gγ2 subunits (Fig 5A). We showed that CXCL12 induced a dose-dependent decrease of the BRET^2^ signal for the three Gαi proteins (Gαi1, Gαi2 and Gαi3) while it did not activate significantly the other G proteins (Fig 5A and S6 Fig). The CXCR4 antagonist AMD3100 and a CXCR4 blocking antibody (MAB173) both reduced the change in BRET^2^ signal, demonstrating its specificity (S7 Fig). Interestingly, coexpression of CCR7 with CXCR4 resulted in a selective inhibition of Gαi1 and Gαi2 activation by CXCL12, while the profile of Gαi3 activation was insensitive to CCR7 (Fig 5A and S6 Fig). As shown for cells expressing CXCR4 only, pretreatment of cells coexpressing CXCR4 and CCR7 with CXCR4-blockers completely inhibited Gαi3 activation, confirming that this signal is dependent on CXCR4 activation (S7 Fig).

**Figure 5.**
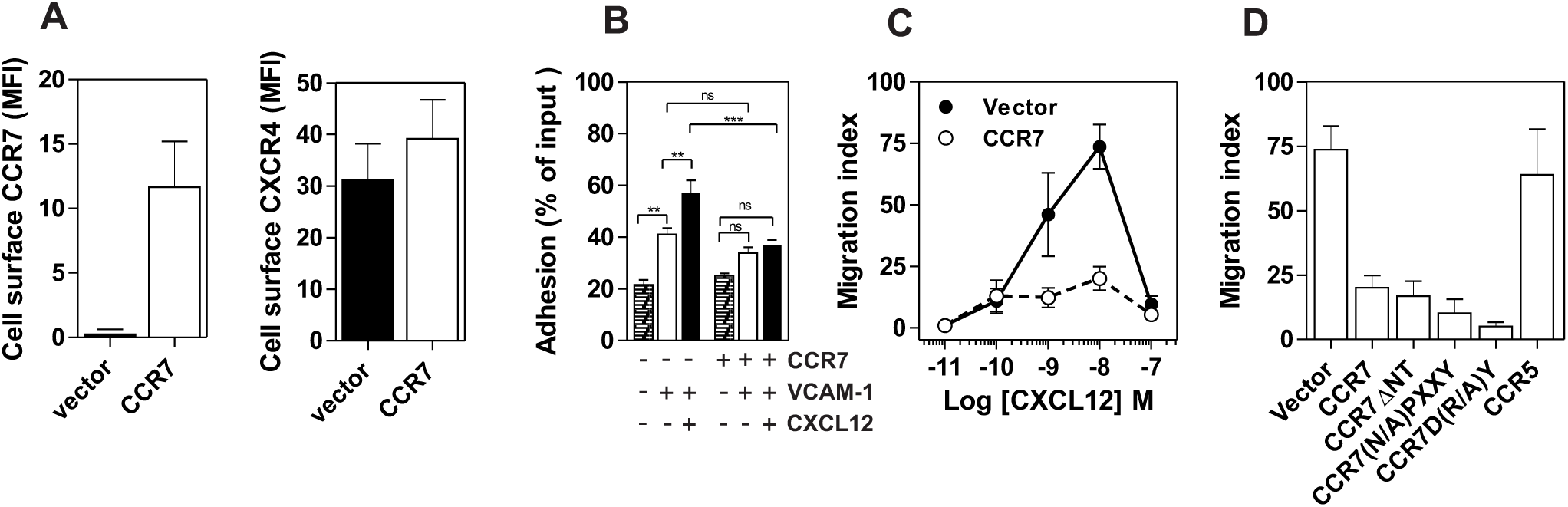
**A. Expression of CCR7 in Nalm-6 does not inhibit CXCR4 expression.** Nalm-6 cells were stably transfected with a CCR7-encoding plasmid and cell surface expression of CXCR4 and CCR7 was monitored by flow cytometry using PE-conjugated anti-CCR7 or anti-CXCR4 antibodies. The data represent mean values of the mean fluorescent index ± SEM (n = 3). **B. Expression of CCR7 in Nalm-6 cells inhibits CXCL12-induced adhesion to VCAM-1.** A suspension of Nalm-6 cells expressing CCR7 or not were stimulated with 1 µM CXCL12 for 1 minute (black bars) or incubated in buffer (white bars) and then allowed to settle in VCAM-1-coated wells for 1 minute. Uncoated wells were used as controls (dashed bars). Non-adherent cells were subsequently washed away and adherent cells were counted relative to the number of input cells. The data represent mean values ± SEM (n = 3; **P < 0.05; P<0.005). **C. Expression of CCR7 in Nalm-6 cells inhibits CXCL12-induced chemotaxis.** Migration of Nalm-6 cells expressing CCR7 (white dots) or not (black dots) was recorded in Transwells in response to increasing concentrations of CXCL12. The data represent mean values ± SEM (n = 3). **D. Expression of non-functional CCR7 mutants also inhibits CXCR4 responsiveness.** Transwell migration of Nalm-6 cells stably expressing wild-type or non-functional CCR7 mutants was assayed in response to 10 nM CXCL12. Cells stably expressing CCR5 were used as control. The data represent mean values ± SEM (n = 3).

### CCR7 modifies the interaction between CXCR4 and G proteins

We next tested whether the lack of Gαi1 and Gαi2 activation was associated with their inability to interact with CXCR4 in the presence of CCR7, and measured the interaction between h*R*Luc-Gαi proteins and CXCR4-Venus by BRET^1^. It was previously demonstrated that BRET^1^ can sense conformational rearrangements within receptor-G protein pre-complexes when measuring the interaction between receptor and G protein subunits [21, 22]. CCR7 expression decreased the basal BRET^1^ signal between *h*RLuc-Gαi1 and CXCR4-Venus or between hRLuc-Gαi2 and CXCR4-Venus (Fig. 5B). In contrast, basal BRET^1^ signal between *h*RLuc-Gαi3 and CXCR4-Venus is less affected by the presence of CCR7 (Fig 5B). A decrease in the BRET signal might indicate that CCR7 impairs the interaction of Gαi1 and Gαi2 proteins with CXCR4. Alternatively, these modifications of BRET could reflect distinct conformations of CXCR4/Gαi protein complexes resulting from CXCR4 heteromerization with CCR7, as previously reported for other CXCR4 heteromers [24, 25].

### CCR7 interacts with CXCR4

We first investigated the ability of CXCR4 to interact with CCR7 by using BRET-proximity assay. A specific BRET^1^ signal was detected between CXCR4-*h*RLuc and CCR7-Venus as well as between CCR7-*h*RLuc and CXCR4-Venus, while a much lower BRET^1^ signal was detected between CXCR4-*h*RLuc and the TSHR-Venus, used as a specificity control (Fig 6A). Further support for physical and direct interactions between CXCR4 and CCR7 came from Bimolecular Fluorescence Complementation experiments, in which each receptor is fused to complementary fragments of the Venus protein (V1 and V2) [7]. Co-expression of CXCR4-V1 with CCR7-V2 generated an important fluorescent signal, confirming the formation of CXCR4/CCR7 heteromers (Fig 6B). A much weaker fluorescence was detected when cells expressed a single chimera or coexpressed the chimera with TSHR-V1 or TSHR-V2. Previously, we showed that heteromerization of CXCR4 with other chemokine receptors results in a strong negative binding cooperativity [6, 7]. Using the same methodology, we showed here that co-expression of CXCR4 and CCR7 is not associated to negative binding cooperativity. CXCL12 did not inhibit the binding of radiolabelled-CCL19 and, conversely, CCL19 and CCL21 did not inhibit the binding of radiolabelled-CXCL12 (Fig 7A and 7B). Nevertheless, the homologous competition performed in cells coexpressing CXCR4 and CCR7 unveiled a second CXCL12 binding site of low affinity (Fig 7A, right panel), which may indicate the presence of a G protein-uncoupled state of CXCR4 within CXCR4/CCR7 heteromers. Importantly, we confirmed that CCR7 inhibits the CXCR4 responsiveness in CHO-K1 cells as measured by calcium mobilization or receptor endocytosis assays (Fig 7C and 7D). Collectively, these results demonstrate that CCR7 interacts with CXCR4 and modifies the conformation of the CXCR4/G protein complexes. It is therefore tempting to link these conformation modifications to the inability of CXCR4 to activate Gαi1 and Gαi2 in the presence of CCR7.

**Figure 6.**
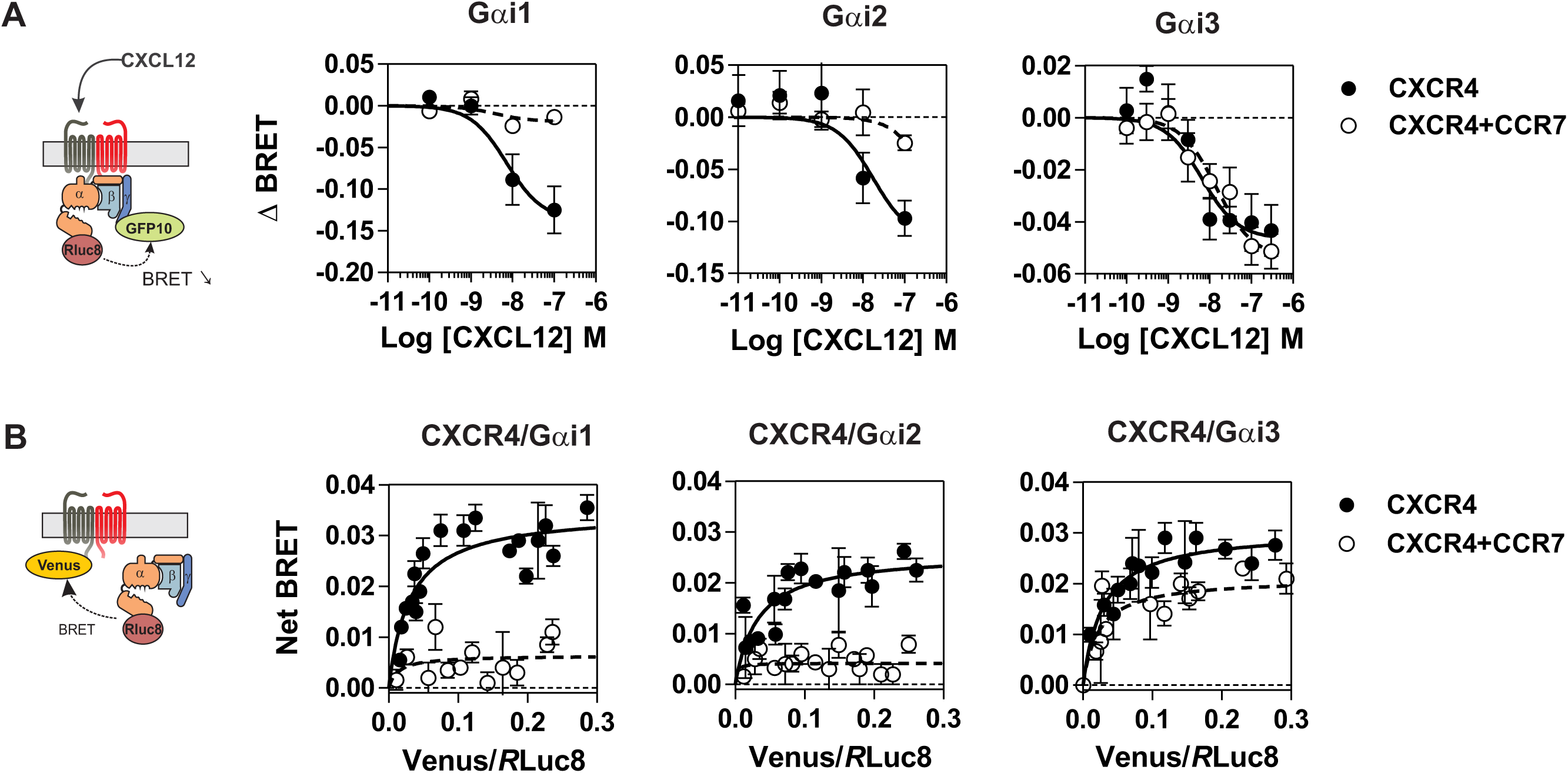
**A. CCR7 inhibits the CXCL12-induced activation of Gαi1 and Gαi2 proteins.** Real-time measurement of BRET signal in HEK293T cells coexpressing Gαi1, Gαi2 or Gαi3 biosensors with CXCR4 only (black dots) or in a combination with CCR7 (white dots). Cells were stimulated for one minute with increasing concentrations of CXCL12 after addition of coelenterazine 400. Results are expressed as ΔBRET, corresponding to the difference in BRET signal between Gαi-*h*RLuc8 and Gβ1γ2-GFP10, measured in the presence and absence of CXCL12. The data represent mean values ± SEM (n = 6). **B. CCR7 changes the basal BRET signal between CXCR4 and G proteins.** HEK293T cells were transfected with h*R*Luc-Gαiβγ and CXCR4-Venus in combination or not with CCR7, and interaction between Gαiβγ and CXCR4 was investigated by measuring the energy transfer (BRET^1^) between the partners. The net BRET corresponds to the BRET measured between the two partners minus the BRET measured in cells expressing h*R*Luc-Gαiβγ only. The data represent mean values ± SEM (n = 5; ***P < 0.0005).

**Figure 7.**
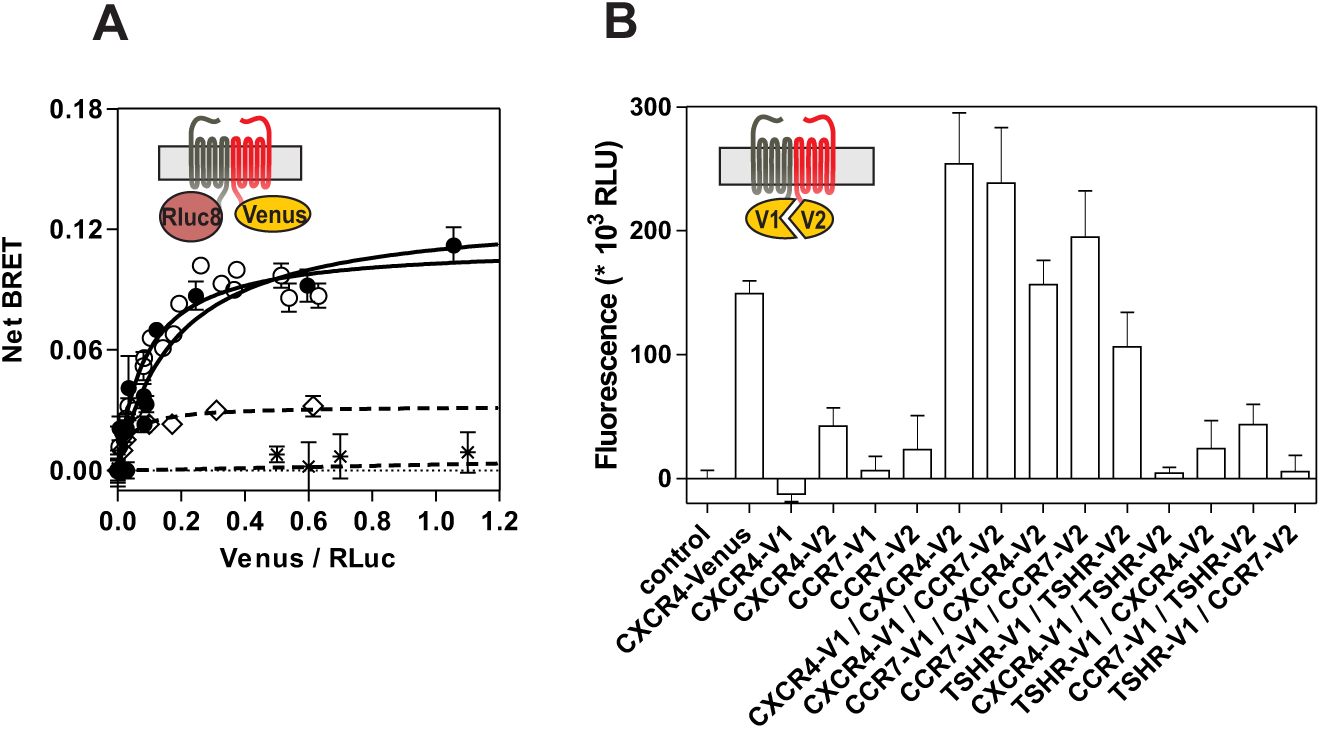
**A. CCR7 interacts with CXCR4 in a BRET assay.** HEK293T cells were transfected with a constant amount of the CCR7-h*R*Luc fusion and increasing amounts of the CXCR4-Venus fusion (black dots), or a constant amount of the CXCR4-h*R*Luc fusion and increasing amounts of the CCR7-Venus fusion (white dots), and heteromerization of CCR7 and CXCR4 was investigated by measuring the energy transfer (BRET^1^) between the two partners. As a control, an increasing amount of TSHR-Venus was used as BRET acceptor with CXCR4-h*R*Luc (*) or CCR7-h*R*Luc (◊) as donor. The Net BRET corresponds to the BRET measured between the two partners minus the BRET measured in cells expressing CXCR4-h*R*Luc or CCR7-h*R*Luc only. Data represent mean values ± SEM (n = 3). **B. CCR7 interacts with CXCR4 in a fluorescence complementation assay.** HEK293T cells were transfected with CXCR4-V1, CXCR4-V2, CCR7-V1 and CCR7-V2 constructs, alone or as two by two combinations, and the fluorescence emission was recorded. As controls, TSHR-V1 and TSHR-V2 were cotransfected with the various CXCR4 and CCR7 constructs. Data represent mean values ± SEM (n = 3).

**Figure 8.**
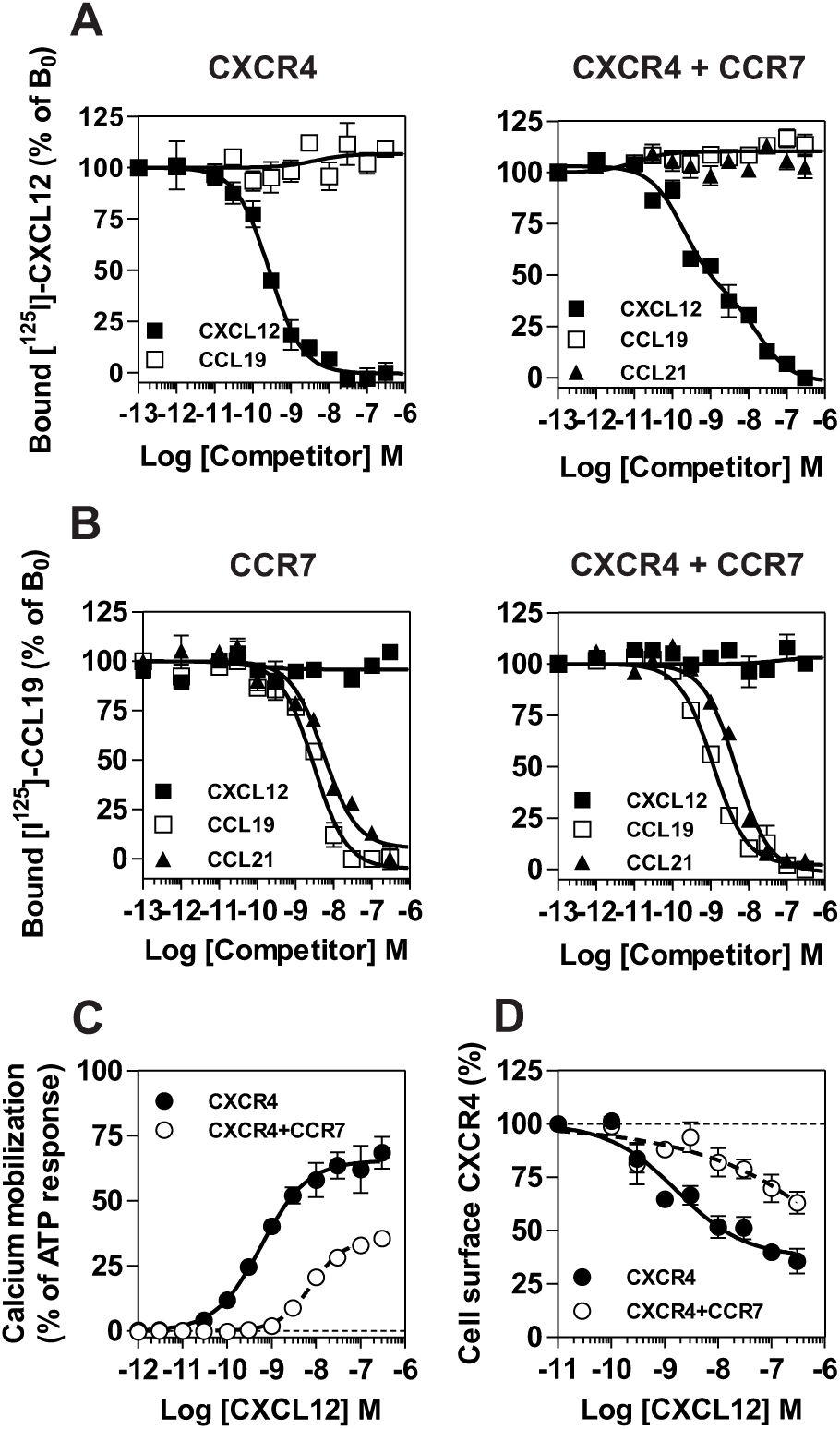
**A-B. Competition binding assays in CHO-K1 cells co-expressing CCR7 and CXCR4.** Competition binding assays were performed on cells expressing CXCR4 or CCR7 only and on cells co-expressing CXCR4 and CCR7. Cells were incubated with 0.1 nM ^125^I-CXCL12 (**A**) or 0,1nM ^125^I-CCL19 (**B**) as tracers and increasing concentrations of unlabeled CXCL12 (black squares) or CCL19 (white squares) as competitors. The data were normalized for nonspecific binding (0%) in the presence of 300 nM of competitor, and specific binding in the absence of competitor (100%). All points were run in triplicates and the data are presented as mean values ± SEM (n = 3). **C. CCR7 inhibits CXCR4 responsiveness in a calcium mobilization assay**. Functional responses of CHO-K1 cells expressing CXCR4 only or in combination with CCR7 were measured using the aequorin-based calcium mobilization assay. Cells were loaded with coelenterazine H, stimulated with increasing concentrations of CXCL12 and the luminescence was recorded. The results were normalized for baseline activity (0%) and the maximal response obtained with 25 µM ATP (100%). All points were run in triplicates and the data are presented as mean values ± SEM (n = 3). **D. CCR7 inhibits the downmodulation of CXCR4 induced by CXCL12**. CHO-K1 cells expressing CXCR4 only or in combination with CCR7 were either left untreated or stimulated for 90 minutes with increasing concentrations of CXCL12. Surface-bound CXCL12 was removed by an acid wash step and cell surface expression CXCR4 was estimated by FACS. The data were normalized for the expression of receptor in absence of stimulation (100%). All points were run in duplicates and the data represent mean values ± SEM (n = 3).

## Discussion

B cells arising from hematopoietic stem cells go through a series of developmental stages in the bone marrow. Soluble factors produced by stromal cells within specific marrow niches, such as the chemokine CXCL12, sustain the development and retention of B cell precursors as they differentiate [26, 27]. Ultimately, about two-thirds of immature B cells are released into the peripheral blood circulation to reach the spleen and complete their maturation, whereas the remaining B cells mature directly in the bone marrow [28, 29]. When bone marrow pre-B cells differentiate into immature and mature B cells, the responsiveness of CXCR4 drops dramatically, despite the fact that high cell surface expression of the receptor is maintained [14]. Such dissociation between CXCR4 expression and responsiveness to CXCL12 has also been reported in other cell types, as well as for other chemokine receptors, but the exact reason for the discrepancy between receptor expression and function often remains undetermined [30,31,32].

In this paper, we show that the cell surface expression of chemokine receptors changes significantly during B cell development in bone marrow. Surface expression of CXCR4 decreases during the transition of pre-B cells to immature and mature B stages, whereas expression of CCR7 and other chemokine receptors increases. In CCR7-deficient mice, CXCR4 responsiveness is improved in mature B cells, and to a lesser extend in immature B cells. The recovered CXCL12-induced migration in CCR7^-/-^ mature B cells is relatively modest compared to the migration of pre-B or immature B cells, though the migration index is similar to that triggered by CCL19 or CXCL13. These results suggest that mature B cells have a relatively weak propensity for chemotaxis. In contrast, when testing calcium mobilization, we showed that the response to CXCL12 is similar for CCR7^-/-^ mature B cells and wild type immature B cells, suggesting that CCR7 regulates CXCR4 responsiveness, but that the relative efficiency of triggering chemotaxis and calcium mobilization varies according to cell type. This hypothesis is in the line with the new paradigm of signaling selectivity. We can’t formally exclude however that other factors than CCR7 might control CXCR4 responsiveness. Analyzing the ImmGen database (www.Immgen.org) reveals that several genes encoding signaling or regulatory proteins are up- or down-regulated between pre-B cells and mature B cells. Another study shows that the expression of genes coding for signaling proteins and cell adhesion molecules is modulated during B cell development [33]. However, it is difficult to appreciate whether these elements contribute to the regulation of CXCR4 responsiveness. Thus, although the precise mechanisms controlling the chemotactic behavior and more particularly the CXCR4 responsiveness during B cell development remain to be identified, our results clearly show that CCR7 expression participates to the regulation of CXCR4 function. A moderate increase of CXCR4 responsiveness is also detected in immature B cells of CCR7^-/-^ mice. One might speculate that additional mechanisms contribute to the regulation of CXCR4 responsiveness and that the impact of CCR7 deficiency is only detectable in certain cellular context.

Labeling cells located in bone marrow sinusoids showed that the proportion of immature and mature B cells located in parenchyma is higher in CCR7^-/-^ mice, while the number of B cells in sinusoids decreases in these mice. These results are in line with the known role of the CXCL12/CXCR4 axis in the homing of B cell precursors in bone marrow and suggest that CCR7 regulates indirectly the distribution of cells between the parenchyma and the sinusoids [34, 19]. Despite a decrease in the egress of immature and mature B cells to the sinusoids of CCR7^-/-^ mice, we showed that the number of these cells is similar in the blood of CCR7^-/-^ and wild type mice. A similar B cell blood count despite a general defect in the distribution of various leukocyte populations amongst different organs and the bloodstream was previously described in CCR7^-/-^ mice [20].

We confirm in adoptive transfer experiments that CCR7-deficient immature B cells home more readily to bone marrow. This homing is strictly dependent on CXCR4 as AMD3100 treatment completely abolished it. In striking contrast, CCR7-deficiency did not affect the recruitment of transplanted mature B cells to bone marrow. These results may appear paradoxical in the light of the results generated in *ex vivo* chemotaxis assays, in which the migration of mature but not immature B cells is enhanced in the absence of CCR7. However, the inability of AMD3100 to affect the number of transplanted mature B cells suggests that CXCL12 is not the predominant signal for the short term homing of mature B cells and that other signals play major roles in the behavior of these cells *in vivo*. In keeping with this hypothesis, signals mediated by CB2 receptor were recently shown to affect bone marrow retention of immature and mature B cells [19]. The migration index of mature B cells appears also relatively low *ex vivo*, suggesting that the CXCL12/CXCR4 pathway might efficiently control their retention in bone marrow while contributing modestly to the recruitment of these cells from the bloodstream.

We also clearly showed that the more efficient retention of B cells in the bone marrow of CCR7^-/-^ mice is not due to an alteration of the bone marrow environment. When we transplanted bone marrow from CCR7^-/-^ mice to irradiated wild type mice, the number of B cells decreased significantly and the relative proportion of the mature B cell subpopulation increased. This result might indicate that CCR7 is directly involved in hematopoiesis as reported for T cells [35]. Alternatively, a higher number of mature B cells in the bone marrow might also affect the hematopoiesis process through feedback inhibition as recently suggested [36]. Whether CCR7 influences additional B cell behaviors than retention in the parenchyma remains thus to be determined.

We also investigated the molecular mechanisms through which CCR7 inhibits CXCR4 responsiveness. First, we showed that the inhibition of CXCR4 by CCR7 is selective as CCR7 is not involved in the regulation of other chemokine receptors, CCR6 and CXCR5, also expressed by mature B cells. This hypothesis is further supported by previous data showing that CCR7 does not inhibit the function of ChemR23 in recombinant cells, although receptor heteromers were formed [37]. Similarly, heteromerization of CXCR4 with other chemokine receptors does not lead to functional inhibition of CXCR4 [6,7,37,38]. It appears therefore that the functional consequences of receptor co-expression and interaction are highly dependent on the specific pair of receptors considered. We also showed that CCR7 signaling is not required, as inhibition of CXCR4 function occurred in the absence of CCR7 stimulation. Moreover, inhibition of CCR7 by blocking antibodies or the use of non-functional CCR7 mutants did not alleviate the inhibition of CXCR4 responsiveness. These findings support a dual role of CCR7 as signaling receptor and modulator of CXCR4. The reduced responsiveness of CXCR4 is neither due to its inability to interact with CXCL12, as saturation binding assays showed that CXCL12 binds as efficiently to CXCR4 whether CCR7 is expressed or not. This observation is hardly compatible with a complete inhibition of interaction with Gαi proteins, known to be required for high affinity binding of chemokines. Nevertheless, in competition binding assays, a second low-affinity binding site for CXCL12 was unveiled in cells coexpressing CXCR4 and CCR7. This result may suggest the existence of two CXCR4 populations, one of which displaying a reduced propensity to interact with CXCL12. The existence of a CXCR4 population with low binding affinity may account at least partly for the decrease of CXCR4 responsiveness detected in native and recombinant cells. It was reported for some GPCRs that heteromerization modifies their binding properties and, by using BRET and functional complementation assays, we confirmed that CXCR4 physically interacts with CCR7. The Gαi proteins interact with both CXCR4 homomers and CXCR4/CCR7 heteromers. An altered conformation of CXCR4/Gαi complex might thus explain the low responsivenes detected in cells co-expressing CXCR4 and CCR7. By using G protein biosensors, we showed that the activation of Gαi1 and Gαi2 proteins by CXCL12 is reduced in cells coexpressing CXCR4 and CCR7. The inhibition of the Gαi2 activation likely accounts for the decrease of B cell retention in bone marrow, as Gαi2 was shown to be required for CXCR4 signaling in the frame of B cell chemotaxis [39, 40]. It is more difficult to appreciate whether the inhibition of Gαi1 also contributes to the regulation of B cell retention as murine lymphocytes express weakly Gαi1 compared to Gαi2 and Gαi3. Our results also suggest that activation of Gαi3 by CXCR4 is maintained in the presence of CCR7. Although this result is potentially interesting, it is difficult to confirm the persistence of signaling, as the nature of the signals downstream of Gαi3 protein are not precisely known [41]. Whether Gαi3-dependent signaling still exists in cells coexpressing CXCR4 and CCR7 remains thus an open question that will require further analysis.

We showed in this study that CXCR4/CCR7 heteromerization results in deficient activation of Gαi1 and Gαi2 proteins. The upregulation of CCR7 and the concomitant decrease of CXCR4 during B cell development might favor the formation of heteromers, leading eventually to a defect in Gαi1/2 signaling. Our data support thus a functional consequence of CXCR4 heteromerization in a physiological process. It has been previously shown that CXCR4 can interact with several other GPCRs, including the chemokine receptors CCR2, CCR5, CXCR7, the Epstein-Barr Virus-encoded BILF1 as well as with the β2 adrenergic receptor. Heteromerization of CXCR4 with CCR2 and CCR5 results in a strong negative binding cooperativity of allosteric nature *i.e.* the specific ligands of CCR2 and CCR5 inhibiting the binding of CXCL12 onto CXCR4 [6, 7]. The coexpression of CXCR7 with CXCR4 was reported to inhibit activation of Gαi proteins by CXCR4 [24]. However, whether this effect requires CXCR4 heteromerization remains to be determined precisely, as CXCR7 can also bind CXCL12 with high affinity and promote a range of cellular responses by itself [42, 43]. In another study, it was reported that coexpression of CXCR7 and CXCR4 results in a constitutive recruitment of β-arrestin 2 to CXCR4/CXCR7 heteromers and enhanced cell migration [44]. Coexpression of BILF1 and CXCR4 was shown to inhibit almost completely CXCL12 binding to CXCR4 as a consequence of BILF1 constitutive activity, but it is unknown whether this effect is linked to CXCR4/BILF1 heteromerization [45]. Finally, β2 adrenergic receptor activation was reported to enhance CXCR4 signaling, promoting the retention of T lymphocytes in lymph nodes, but whether this increase in CXCR4 responsiveness is caused by its physical interaction with β2 remains to be established [46].

Our findings reveal thus the existence of a novel mechanism of regulation of CXCR4 function and they constitute, to our knowledge, the first indication that chemokine receptors coexpression has consequences on a physiological process. The regulation of CCR7 expression might also affect CXCR4 responsiveness in other cell types, such as T cells, macrophages or dendritic cells. Even if testing this hypothesis is beyond the scope of the current study, we have to take into account in future investigations that the functional consequence of CCR7 expression on CXCR4 function might also vary according to the cellular context. In contrast to the situation encountered with mature B cells isolated from bone marrow, mature B cells isolated from spleen migrate very efficiently toward CXCL12, and the consequence of CCR7 deficiency on splenic cell migration is almost undetectable. Whether CCR7 controls CXCR4 functions differently according to cell types thus remains to be determined precisely.

Our results may also have important implications in pathological contexts. Several studies have highlighted the critical role of the CXCL12-CXCR4 axis in cancer progression, and overexpression of CXCR4 by tumor cells is generally considered as a poor prognosis marker. Yet, a lack of correlation between CXCR4 expression and cell migration has been reported in some studies [47, 48]. The expression of accessory proteins like ZAP70 or CXCR7 in tumors was shown to regulate the function of CXCR4 [49, 50]. Heteromerization of CXCR4 and CCR7 in some tumors may be an alternative mechanism to explain the lack of correlation between expression of CXCR4 and its responsiveness. Our discovery that CXCR4 function in vivo depends on the expression of another chemokine receptor adds thus a new dimension to our understanding of how CXCR4 might control cell behavior in physiological and pathological processes.

## Material and methods

### Mouse models

The CCR7-deficient mouse line, described previously [20], was obtained from The Jackson Laboratory and backcrossed on the C57BL/6 background for at least 8 generations. The Boy/J (CD45.1^+^) mouse line was obtained from Janvier laboratory. All mice were housed under specific-pathogen-free conditions at our local mouse facility. In all experiments, CCR7-deficient mice (6 to 8 weeks old) were compared with wild-type littermates resulting from CCR7^+/-^ intercrossing. To generate bone marrow chimeras, recipient mice were irradiated lethally (10 Gy) and their immune system was reconstituted by intravenous injection of 2 10^6^ total bone marrow cells from donor mice. Chimera were analyzed 8 to 10 weeks after reconstitution. Animal experimentation was carried out in accordance with European (EU Directives 86/609/EEC) and national guidelines. All procedures were reviewed and approved by the local ethical committee (Commission d’Ethique du Bien-Etre Animal, CEBEA) of the Université Libre de Bruxelles.

### Expression of recombinant receptors and mutants

All the receptors used in this study were either cloned into bicistronic pEFIN3 vector (Euroscreen) for stable expression or pcDNA3 vector (Life Technologies) for transient expression (S1 Table). All constructs were verified by sequencing prior to transfection.

### Cell culture and transfections

The human pre-B cell line Nalm-6 (ACC-128, DMSZ) was cultured in RPMI 1640 containing L-glutamine supplemented with 10% heat-inactivated FBS, 100 U/ml penicillin, 100 µg/ml streptomycin and 50 mM 2-mercaptoethanol (Sigma). Transfection of Nalm-6 cells by pEFIN3 plasmids was performed by electroporation (0.3 kV, 960 µF) and cells stably expressing the receptors of interest were selected and cultured in the presence of 800 µg/ml G418 (Invitrogen). The human embryonic kidney cell line HEK293T (CRL-3216, ATCC) was cultured in

Dulbecco’s Modified Eagle’s Medium (DMEM) supplemented with 10% FBS (GIBCO), 100 U/ml penicillin and 100 µg/ml streptomycin (Invitrogen). Transfection of HEK293T cells by pcDNA3 plasmids was performed 24 h after cell seeding using the calcium phosphate precipitation method.

### Flow cytometry

Mononuclear cells from blood, bone marrow, spleen or lymph nodes were isolated by Ficoll-Paque separation, diluted in staining buffer (PBS with 0.1% BSA and 0.01% sodium azide) and incubated with 1 µg rat anti-mouse FcγR antibody (Clone 2.4G2, BD Biosciences) to prevent binding of conjugated antibodies to FcγR. Cells were further incubated for an additional 30 minutes on ice with a mixture of the following antibodies: Alexa Fluor 700 rat anti-mouse CD45R (B220, clone RA3-6B2, BD Biosciences), FITC rat anti-mouse CD43 (Clone S7, BD Biosciences), PerCP-Cy5.5 rat anti-mouse IgM (clone R6-60.2, BD Biosciences) and V450 rat anti-mouse IgD (clone 11-26c.2a, BD Biosciences). This staining strategy allowed to distinguish the developmental subsets of B cells, namely pre-B cells (B220^+^/CD43^-^/IgM^-^/IgD^-^), immature B cells (B220^+^/CD43^-^/IgM^+^/IgD^-^) and mature B cells (B220^+^/CD43^-^/IgM^+^/IgD^+^) [17, 18]. As an alternative staining strategy, cells were incubated with a mixture of Alexa Fluor 700 rat anti-mouse CD45R, PerCP-Cy5.5 rat anti-mouse IgM, PE rat anti-mouse AA4.1 (clone AA4.1, BD Biosciences) and FITC rat anti-mouse CD24/HSA (clone M1/69, BD Biosciences) to discriminate pre-B cells (B220^+^/IgM^-^/AA4.1^+^/HSA^+^), immature B cells (B220^+^/IgM^+^/ AA4.1^+^/HSA^+^) and mature B cells (B220^+^/CD43^+^/ AA4.1^low^/HSA^low^) [51]. Absolute cell numbers were determined by incorporating beads in the cell suspension (15 µm, Bangs Laboratories) and acquiring 15,000 beads. Expression of chemokine receptors on B cell subsets was estimated by using the following antibodies: phycoerythrin anti-mouse CXCR4 (clone 2B11, eBioscience), allophycocyanin anti-mouse CCR7 (clone 4B12, eBioscience), allophycocyanin anti-mouse CXCR5 (clone 614641, R&D Systems) and allophycocyanin rat anti-mouse CCR6 (clone 140706, R&D Systems).

### RNA extraction and RT-qPCR

RNA was extracted from sorted B cells subpopulations using the RNeasy kit (Qiagen) according to the manufacturer’s instructions. Quantification of gene expression was performed by using the KAPA SYBR-FAST One-Step qRT-PCR kit and the CFX-Connect Real-time PCR detection system (Bio-Rad Laboratories) following the manufacturer’s instructions. The following primers were used: CXCR4 (sense: 5’-AAGAAGTGGGTTCTGGAGAC-3’, anti-sense: 5’-GACTATGCCAGTCAAGAAG-3’), CCR7 (sense: 5’-CCTGCCTCTCATGTATTCTG-3’, anti-sense: 5’-GGTTGAGCAGGTAGGTATCC-3’), CXCR5 (sense: 5’-ATTTTCTTCCTCTGCTGGTC-3’, anti-sense 5’-GAATTCACACAAGGTGATGG-3’), CCR6 (sense: 5’-AGATCATGAAGGATGTGTGG-3’, anti-sense: 5’-TACATGGTAAAGGACGATGC-3’), Beta-actin (sense: 5’-CAGCTTCTTTGCAGCTCCTT-3’, anti-sense: 5’-CACGATGGAGGGGAATACAG-3’) and GAPDH (sense: 5’-AAGGGCTCATGATGACCACAGTC-3’, anti-sense: 5’CAGGGATGATGTTCTGGGCA-3’).

### Chemotaxis assay

The migration of splenic and bone marrow B cell populations in response to chemokine gradients was performed in 6-well Costar transwell chambers (5 µm pore size, Corning). Briefly, a suspension of 10^6^ mononuclear cells in RPMI 1640 supplemented with 10% FCS was added to each insert in a well containing a solution of chemokine (R&D Systems). In some experiments, blocking anti-CXCR4 MAB21651 (247506, R&D Systems) or anti-CCR7 MAB3477 (4B12, R&D Systems) antibodies were added to the cells prior to migration. Wells containing medium without chemokines were used as controls. After 2 h at 37°C, cells in the bottom of the wells were harvested, diluted in FACS staining buffer and incubated with antibodies and counting beads to discriminate and quantify B cell subsets. The migration of Nalm-6 cells was performed in 96-well Costar transwell chambers (5 µm pore size, Corning NY). A cell suspension of 10^4^ Nalm-6 in RPMI 1640 supplemented with 10% FCS was added to each insert. Wells containing medium without chemokines were used as controls. After 1 h at 37°C, cells in the bottom of the wells were counted by using the ATPlite luminescence assay kit (PerkinElmer, Waltham, MA). The results are expressed as chemotaxis index, i.e the ratio of cells migrating in response to the chemoattractant over cells migrating toward the medium alone.

### Calcium mobilization assay

A suspension of 2 10^6^ mononuclear cells was first incubated in PBS with a mixture of antibodies to discriminate B cell subpopulations. Cells were washed and loaded with 4 µM eFluor 514 for 30 minutes at 37°C in RPMI supplemented with 10% FCS. Cells were washed and incubated for 5 minutes at 37°C in 500 µl of RMPI supplemented with 10% FBS before flow cytometry analysis. Fluorescence was recorded for 30 seconds and calcium mobilization was initiated by adding 20 nM CXCL12. After one minute, 5 µg/ml ionomycin was added and the fluorescence was recorded for one more minute. Calcium mobilization was measured in CHO-K1 cells stably expressing chemokine receptors as previously described [7]

### Cell adhesion assay

Short-term adhesion assays were performed as previously described [10]. Briefly, 2.10^4^ Nalm-6 cells in suspension in PBS supplemented with 0.1% BSA were stimulated with 300 nM CXCL12 for one minute and then added to VCAM-1-coated wells (with a 1 mg/ml solution). Cells were quickly spun down and allowed to settle for another minute at 37°C. As controls, cells unstimulated with CXCL12 and uncoated wells were used. Wells were then washed twice and the number of adherent cells was determined using the ATPlite luminescence assay kit (PerkinElmer, Waltham, MA).

### B cell homing assay

B220^+^ B cells were recovered from bone marrow mononuclear cells from WT or CCR7^−/−^ mice by negative selection using a MACS microbeads isolation kit (Miltenyi Biotec) according to the manufacturer’s instructions. Then, 5.10^6^ cells were incubated for 15 minutes in PBS containing 5 µM CFDA succinimidyl ester (Molecular Probes). After washing, about 2.10^6^ CFDA-stained B220^+^ cells were injected retro-orbitally into 6-to 8-week-old syngenic C57BL/6 recipient mice. The percentage of CFDA-labeled cells in the cell mixture was determined by flow cytometry before injection to know the number of CFDA-labelled cells that are transferred. Recipient mice were killed two hours after injection and bone marrow B cell populations (CFDA-labeled or not) were analyzed by FACS.

#### Bi-molecular complementation assay

Bi-molecular fluorescence and luminescence complementation assays were performed in HEK293T cells as described previously [7]. Briefly, plasmids expressing the various receptors fused to split mVenus or Rluc8 fragments were transfected into HEK293T cells. A control corresponding to mock-transfected cells was included in order to subtract raw basal luminescence and fluorescence from the data. Forty-eight hours after transfection, cells were washed twice with PBS, detached and resuspended in PBS. Approximately 2.10^5^ cells were distributed per well of 96-well plates. The 535 nm fluorescence following excitation at 485 nm, and the luminescence after incubation with 5 µM coelenterazine H (Promega) were recorded using a Mithras LB940 reader (Berthold).

#### BRET proximity assay

BRET proximity assays were performed as described previously [6]. Briefly, HEK-293T cells were transfected using a constant amount of plasmid DNA but various ratios of plasmids encoding the fusion protein partners. A control corresponding to mock-transfected cells was included in order to subtract raw basal luminescence or fluorescence from the data. Expression of mVenus fusion proteins was estimated by measuring fluorescence at 535 nm following excitation at 485 nm. Expression of RLuc fusion proteins was estimated by measuring the luminescence of the cells after incubation with 5 µM coelenterazine H (Promega). In parallel, BRET^1^ between h*R*Luc8 and mVenus was measured 5 min after addition of 5 µM coelenterazine H (Promega). BRET^1^ readings were collected using a Mithras LB940 reader (Berthold). The BRET^1^ signal was calculated as the ratio of emission of mVenus (510–590 nm) to h*R*Luc8 (440–500 nm) – Cf, where Cf corresponds to the ratio of emission (510–590 nm) to (440–500 nm) for the hRLuc construct expressed alone in the same experiment.

#### G protein BRET assay

G protein activation was assayed by BRET as previously described [22]. Briefly, plasmids encoding G protein biosensors and receptors of interest were cotransfected into HEK293T cells. Forty-eight hours after transfection, cells were washed twice with PBS, detached and resuspended in PBS containing 0.1% (w/v) glucose at room temperature. Cells were then distributed (80 µg of proteins per well) in a 96-well microplate (Optiplate, PerkinElmer). BRET^2^ between *R*Luc8 and GFP10 was measured 1 min after addition of 5 µM coelenterazine 400a/Deep blue C (Gentaur). BRET readings were collected using an Infinite F200 reader (Tecan). The BRET signal was calculated as the ratio of emission of GFP10 (510–540 nm) to *R*Luc8 (370–450 nm).

#### Statistical analyses

Results are expressed as arithmetic means ± SEM. Significance was determined using Tukey’s test and the Prism4 software (GraphPad). For all tests, values of p<0.05 were considered as significant.

## Acknowledgments

We thank Christine Dubois for her help with cell sorting. S. Mcheik is the recipient of a Télévie studentship. This work was supported by the Fonds de la Recherche Scientifique Médicale of Belgium, the Actions de Recherche Concertées, and the Interuniversity Attraction Poles Programme (P7–40), Belgian State, Belgian Science Policy.

## Supplementary Materials

**Table S1.** Plasmids used in this study.

| Plasmids | Modifications | Position | Reference |
| --- | --- | --- | --- |
| pcDNA3-CXCR4 | - | - | This study |
| pcDNA3-CCR7 | - | - | This study |
| pcDNA3-CXCR4-Ruc8 | RLuc8 | CT | Sohy et al., 2009 |
| pcDNA3-CXCR4-Venus | Venus | CT | Sohy et al., 2009 |
| pcDNA4-CCR7-RLuc8 | RLuc8 | CT | This study |
| pcDNA3-CCR7-Venus | Venus | CT | This study |
| pcDNA3-TSHR-Venus | Venus | CT | This study |
| pcDNA3-CXCR4-V1 | Venus 1-156 | CT | Sohy et al., 2009 |
| pcDNA3-CXCR4-V2 | Venus 157-239 | CT | Sohy et al., 2009 |
| pcDNA3-CCR7-V1 | Venus 1-156 | CT | This study |
| pcDNA3-CCR7-V2 | Venus 157-239 | CT | This study |
| pcDNA3-TSHR-V1 | Venus 1-156 | CT | Sohy et al., 2009 |
| pcDNA3-TSHRV2 | Venus 157-239 | CT | Sohy et al., 2009 |
| pcDNA3-Gαi1 | - | - | cDNA Resource Center (www.cDNA.org) |
| pcDNA3-Gαi2 | - | - | cDNA Resource Center (www.cDNA.org) |
| pcDNA3-Gαi3 | - | - | cDNA Resource Center (www.cDNA.org) |
| pcDNA3-Gβ1 | - | - | Saulière et al., 2012 |
| pcDNA3-γ2 | - | - | cDNA Resource Center (www.cDNA.org) |
| pcDNA3-Gαi1-RLuc8 | RLuc8 | 91-92 | Saulière et al., 2012 |
| pcDNA3-Gαi2-RLuc8 | RLuc8 | 91-92 | Saulière et al., 2012 |
| pcDNA3-Gαi3-RLuc8 | RLuc8 | 91-92 | Saulière et al., 2012 |
| pcDNA3-Gαoa-RLuc8 | RLuc8 | 91-92 | Saulière et al., 2012 |
| pcDNA3-Gαob-RLuc8 | RLuc8 | 91-92 | Saulière et al., 2012 |
| pcDNA3-Gαs-RLuc8 | RLuc8 | 113-114 | Saulière et al., 2012 |
| pcDNA3-Gαq-RLuc8 | RLuc8 | 97-98 | Saulière et al., 2012 |
| pcDNA3-Gα11-RLuc8 | RLuc8 | 97-98 | Saulière et al., 2012 |
| pcDNA3-Gα12-RLuc8 | RLuc8 | 115-116 | Saulière et al., 2012 |
| pcDNA3-Gα13-RLuc8 | RLuc8 | 106-107 | Saulière et al., 2012 |
| pcDNA3-Gγ2-GFP10 | GFP10 | NT | Saulière et al., 2012 |
| pEFIN3-CCR7 | - | - | De Poorter et al., 2013 |
| pEFIN3-HACCR7ΔNT | (1) HA-Tag<br>(2) D/A and E/A | (1) NT<br>(2) D2; D26; E27;D30-31;<br>D35; D40; E45; D52 | This study |
| pEFIN3-CCR7(N/A)PXXY | N/A | N322 | This study |
| pEFIN3-CCR7D(R/A)Y | R/A | R154 | This study |
| pEFIN3-CCR5 | - | - | El-Asmar et al., 2005 |
| NT : N-terminus ; CT :C-terminus |  |  |  |

**Table S2.** Binding parameters of CHO-K1 cells expressing CXCR4 and CCR7.

| Table S2, Binding parameters of CHO-K1 cells expressing CXCR4 and CCR7 |  |  |  |  |
| --- | --- | --- | --- | --- |
| Cells | Tracer | Competitor | IC<sub>50</sub> (nM) | % of inhibition |
| CXCR4 | [<sup>125</sup>I]-CXCL12 | CXCL12 | 0.80 ± 0.05 | 100 |
| CCR7 | [<sup>125</sup>I]-CCL19 | CCL19 | 1.84 ± 1.18 | 100 |
|  |  | CCL21 | 4.06 ± 1.68 | 96.5 ± 6.5 |
| CXCR4<br>+<br>CCR7 | [<sup>125</sup>I]-CXCL12 | CXCL12 | (a) 0.19 ± 0.01<br>(b) 30.88 ± 12.52 | 100 |
|  |  | CCL19 | n.d. | n.d. |
|  |  | CCL21 | n.d. | n.d. |
|  | [<sup>125</sup>I]-CCL19 | CXCL12 | n.d. | n.d. |
|  |  | CCL19 | 1.70 ± 0.46 | 100 |
|  |  | CCL21 | 4.20 ± 0.50 | 98.5 ± 3.5 |
| Binding parameters were measured on cells expressing CXCR4 and CCR7. The IC<sub>50</sub> and % of inhibition values were obtained from competition binding experiments as displayed in Fig. 8. Values represent the mean ± S.E.M. of at least three independent experiments. n.d. not detected. |  |  |  |  |

**Figure S1.**
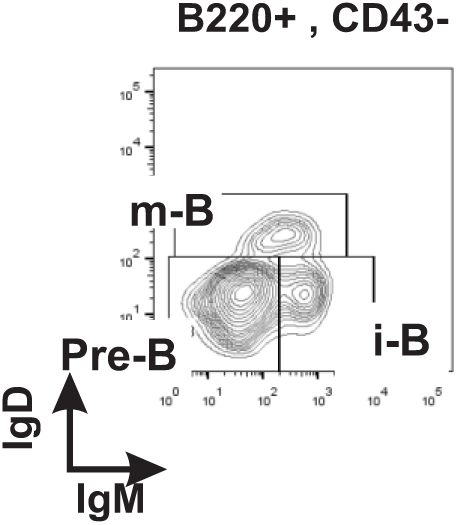
Identification of B cell subpopulations. Bone marrow B cell subpopulations were discriminated by flow cytometry using AF700-conjugated anti-B220, FITC-conjugated anti-CD43, PerCP-Cy5.5-conjugated anti-IgM and V450-conjugated anti-IgD antibodies. Pro/Pre-B cells were identified as B220^+^/CD43^-^/IgM^-^/IgD^-^, immature B cells as B220^+^/CD43^-^/IgM^+^/IgD^-^ and mature B cells as B220^+^/CD43^-^/IgM^+^/IgD^+^. A density plot representative of B cell populations from of a CCR7^+/+^ mouse is shown.

**Figure S2.**
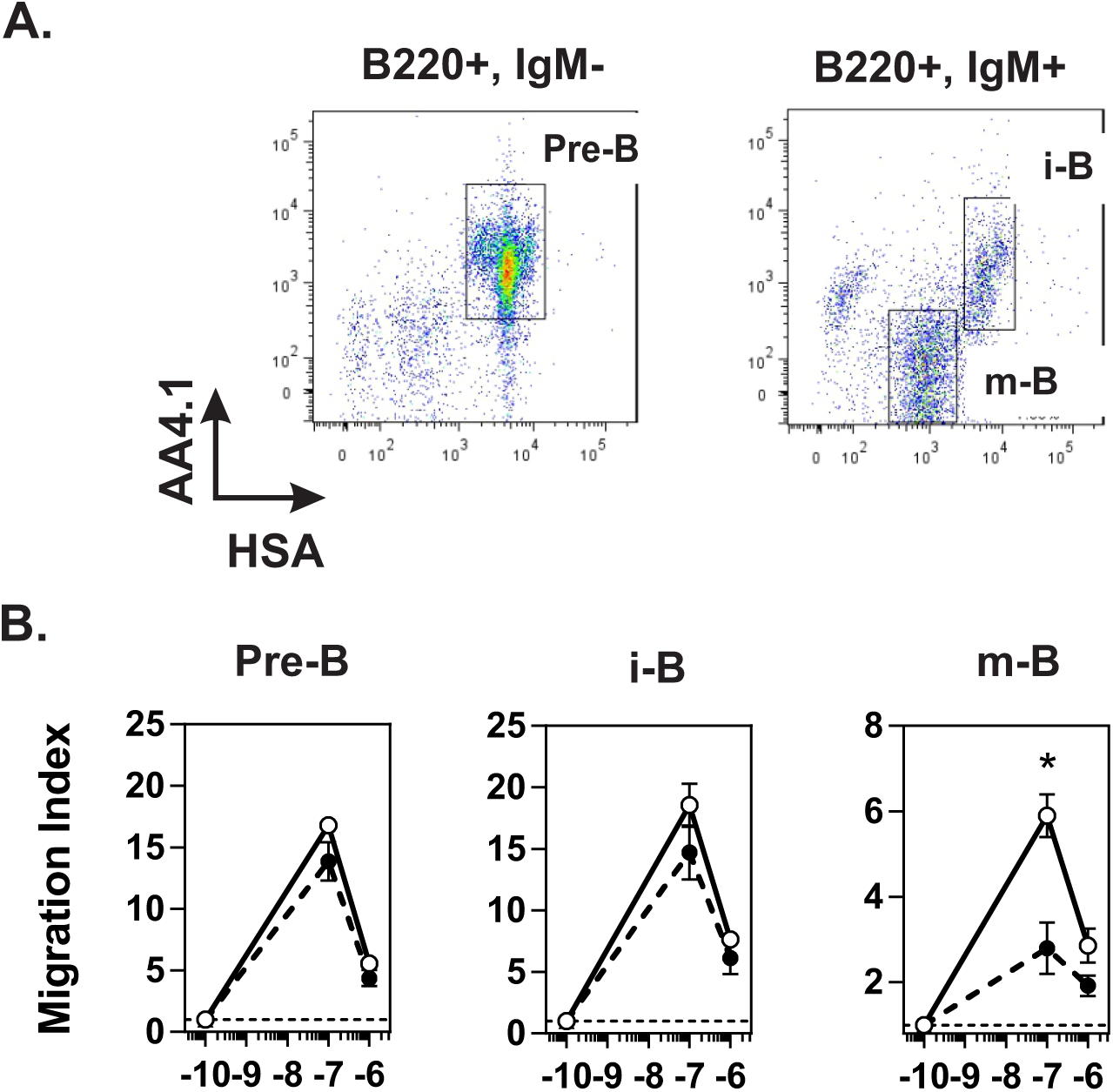
**A. Alternative staining procedure to discriminate B cell subsets.** Bone marrow B cells were discriminated by flow cytometry using AF700-conjugated anti-B220, PercP-Cy5/5-conjugated IgM, PE-conjugated AA4.1 and FITC-conjugated HSA (CD24) antibodies. Pre-B cells were identified as B220^+^, IgM^-^, AA4.1^+^, HSA^+^, immature B cells as B220^+^, IgM^+^, AA4.1^+^, HSA^+^ and mature B cells as B220^+^, IgM^+^, AA4.1 low, HSA low. **B. Chemotaxis of B cells subsets.** Transwell migration of bone marrow cells from CCR7^+/+^ (black dots) or CCR7^-/-^ (white dots) mice in response to 100nM an 1µM CXCL12. Migration index after a 2-h incubation were plotted for Pre-B (B220^+^, IgM^-^, AA4.1^+^, HSA^+^), immature B cells (B220^+^, IgM^+^, AA4.1^+^, HSA^+^) and mature B cells (B220^+^, IgM^+^, AA4.1^low^, HSA^low^). All conditions were run in triplicated and the data are presented as mean ± SEM, * P<0.05.

**Figure S3.**
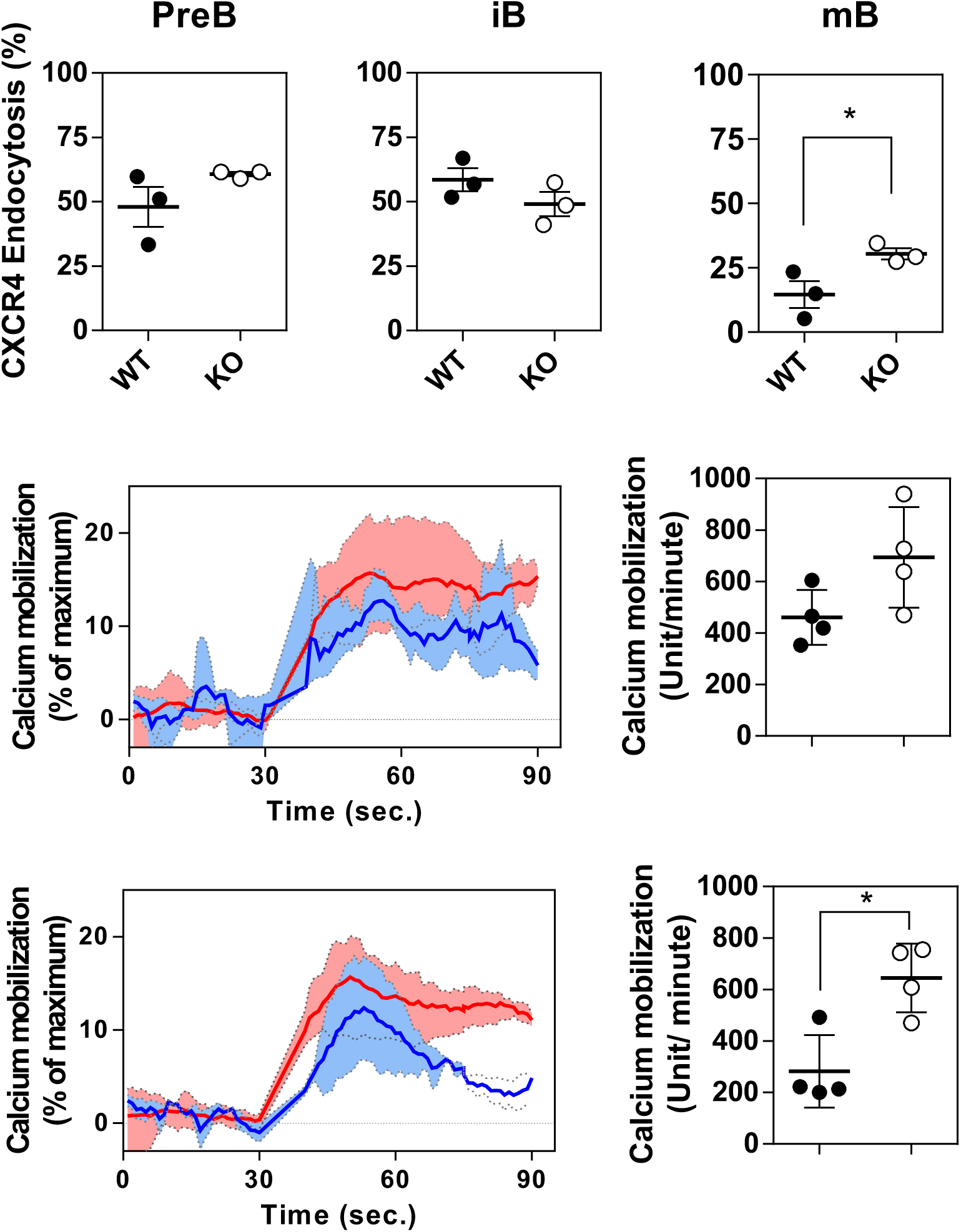
**A. CXCR4 endocytosis in B cell subpopulations.** Bone marrow mononuclear cells were incubated with 50 nM CXCL12 for 45 minutes at 37°C. After an acid wash procedure, cell surface expression of CXCR4 was determined in B cell subpopulations by flow cytometry using AF700-conjugated anti-B220, FITC-conjugated anti-CD43, PerCP-Cy5.5-conjugated anti-IgM, V450-conjugated anti-IgD antibodies and PE-conjugated anti-CXCR4. Data were normalized for cell surface CXCR4 expression in the absence of stimulation (100%) and cell surface fluorescence using PE-conjugated control isotype (0%) and. Results represent the mean ± SEM of three independent experiments were each dot corresponds to individual mice (n= 3, * P<0.05) **B. Calcium mobilization in B cells subsets.** Bone marrow mononuclear cells were first incubated with AF700-conjugated anti-B220, FITC-conjugated anti-CD43, PE-conjugated anti-IgM and V450-conjugated anti-IgD antibodies, for 60 minutes at 4°C, then loaded with eFluor 514 in RMPI for 30 minutes at 37°C. After wash, cells were warmed up at 37°C for 5 minutes and subject to flow cytometry analysis. Thirty second later, calcium mobilization was initiated by adding 20 nM CXCL12. Data were normalized for the fluorescence in the absence of stimulation (0%) and fluorescence in the presence of 5 µg/ml ionomycin (100%). Results represent the mean ± SEM of four independent experiments run in duplicate were each dot corresponds to individual mice (n= 4, * P<0.05)

**Figure S4.**
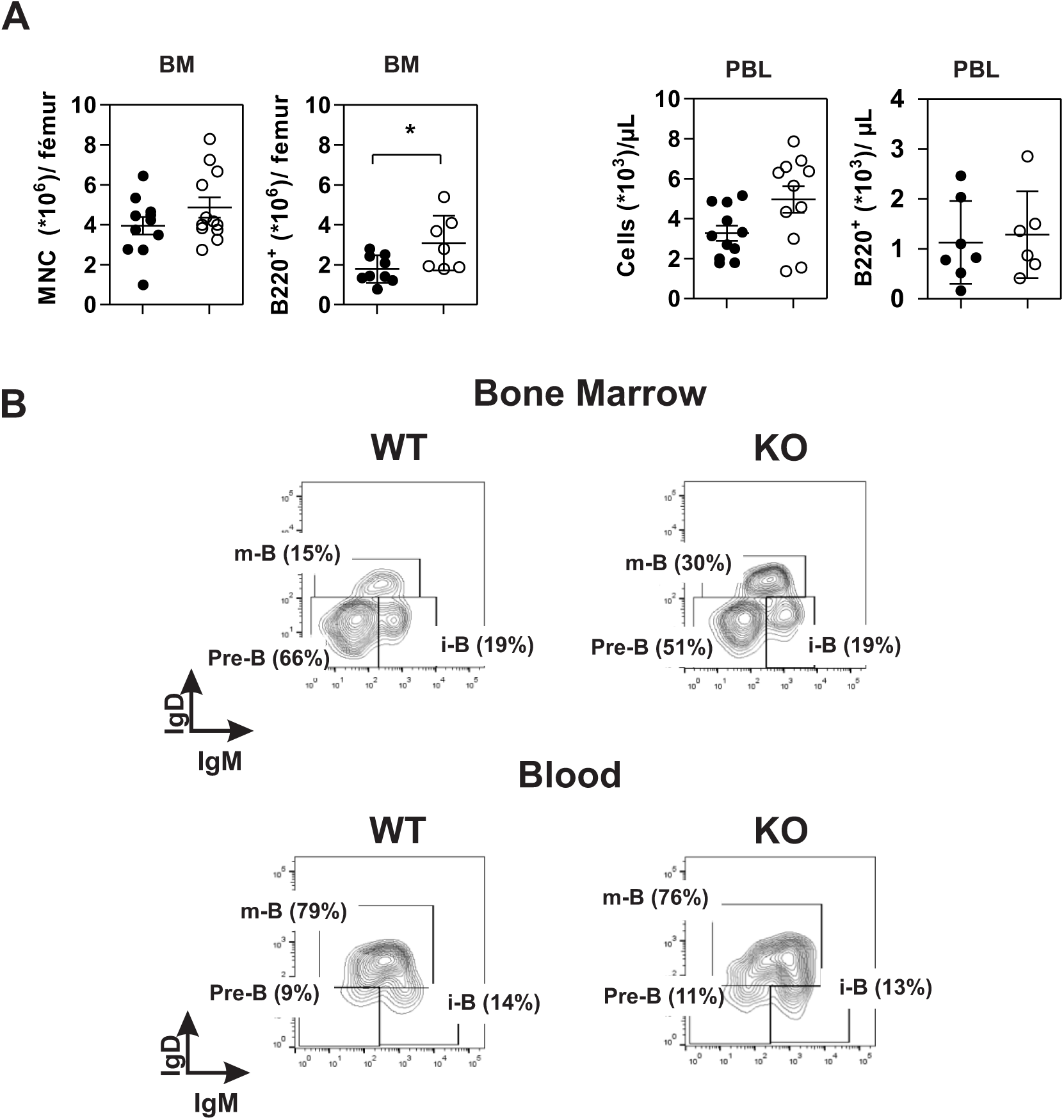
B cells in bone marrow and blood. **A.** Increase number of B cells in the bone marrow of CCR7^-/-^ mice. Number of mononuclear (MCN) and B (B220^+^) cells in the bone marrow of wild type (black dots) and CCR7^-/-^ mice (white dots). Data are represented as the mean values ± SEM and dots corresponds to individual mice (n= 10-13 mice, *, P<0.05). **B.** Representative flow cytometry plots of B cell subpopulations in bone marrow and blood of CCR7^+/+^ and CCR7^-/-^ mice. Pre-B cells (B220^+^, CD43^-^, IgM^-^, IgD^-^), immature B cells as (B220^+^, CD43^-^, IgM^+^, IgD^-^) and mature B cells (B220^+^, CD43^-^, IgM^+^, IgD^+^). Of note, caution should be taken regarding the proportion of Pre-B and i-B in blood as this labelling strategy poorly discriminates these populations.

**Figure S5.**
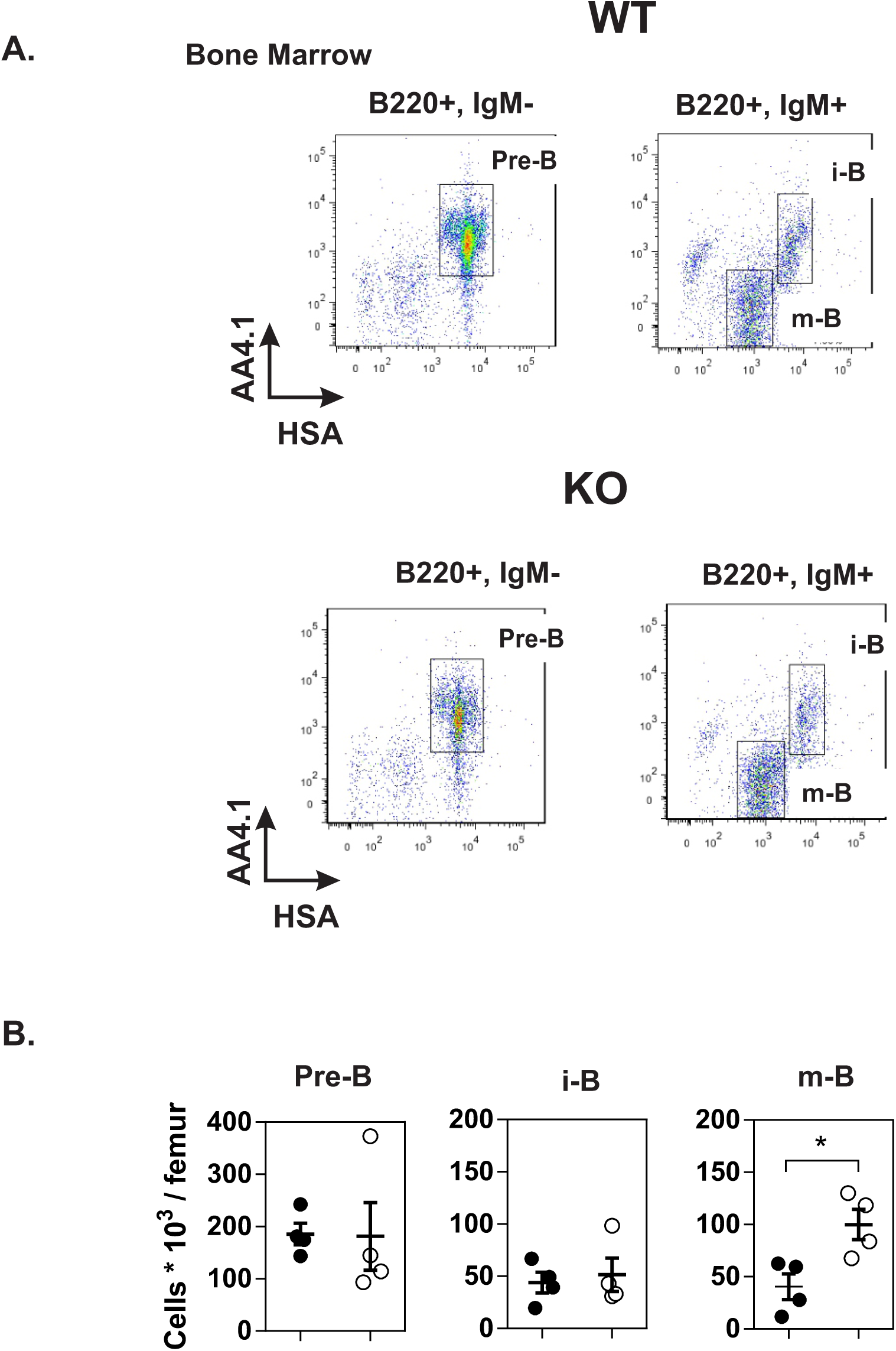
A. B cell subsets in bone marrow identified by the alternative staining procedure. **A.** Representative flow cytometry plots of bone marrow B cell subpopulations of CCR7^+/+^ and CCR7^-/-^ mice. Pre-B (B220^+^, IgM^-^, AA4.1^+^, HSA^+^), immature B cells (B220^+^, IgM^+^, AA4.1^+^, HSA^+^) and mature B cells (B220^+^, IgM^+^, AA4.1 ^low^, HSA^low^). **B. Number of B cells in the bone marrow.** The number of B cell subpopulations in the bone marrow of CCR7^+/+^ (black dots) or CCR7^-/-^ (white dots) mice was analyzed by flow cytometry. Data are presented as mean value ± SEM and dots correspond to individual mice (n=4 mice, * P<0.05).

**Figure S6.**
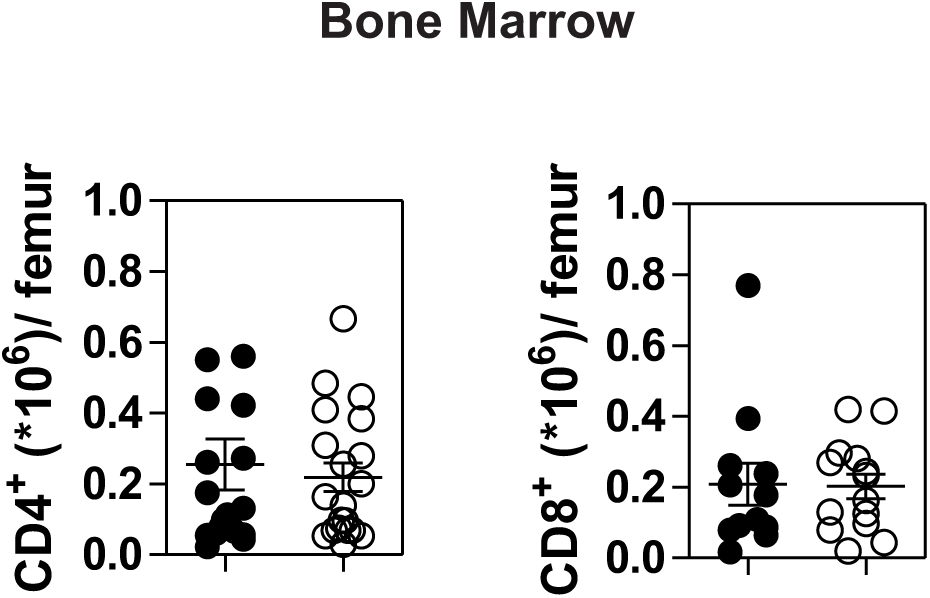
CCR7 deficiency does not increase the amount of T cells in bone marrow. CD4^+^ and CD8^+^ T cells from the bone marrow of CCR7^+/+^ (black dots) and CCR7^-/-^ (white dots) mice were analyzed by flow cytometry. Data are represented as mean values ± SEM where each dot represents individual mice (n= 9 to 16 mice per group).

**Figure S7.**
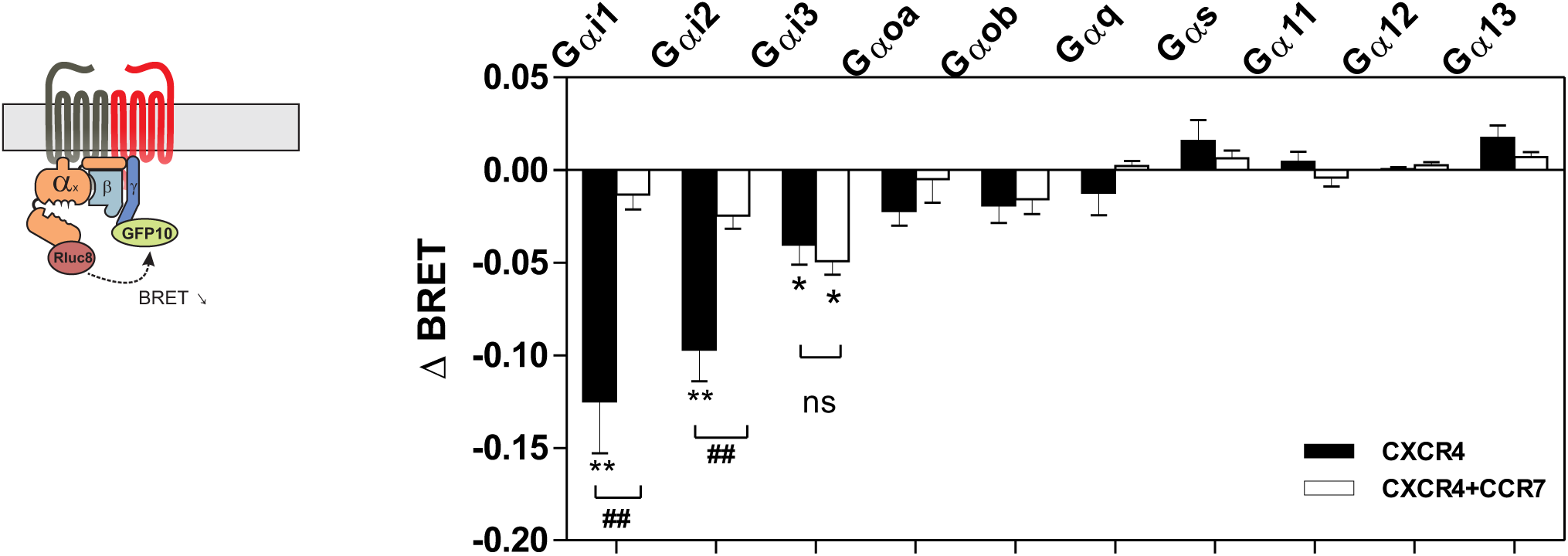
Panel of Gα proteins activated by CXCR4. Real-time measurement of BRET signal in HEK293T cells coexpressing various Gα biosensors with CXCR4 only (black bars) or in combination with CCR7 (white bars). Cells were stimulated one minute with 100 nM CXCL12 after addition of coelenterazine 400. Results are expressed as ΔBRET corresponding to the difference in BRET signal between Gαi-*h*RLuc8 and Gβ1γ2-GFP10 measured in the presence and absence of CXCL12. Data are represented as mean values ± SEM (n= 6, *** P<0,0005).

**Figure S8.**
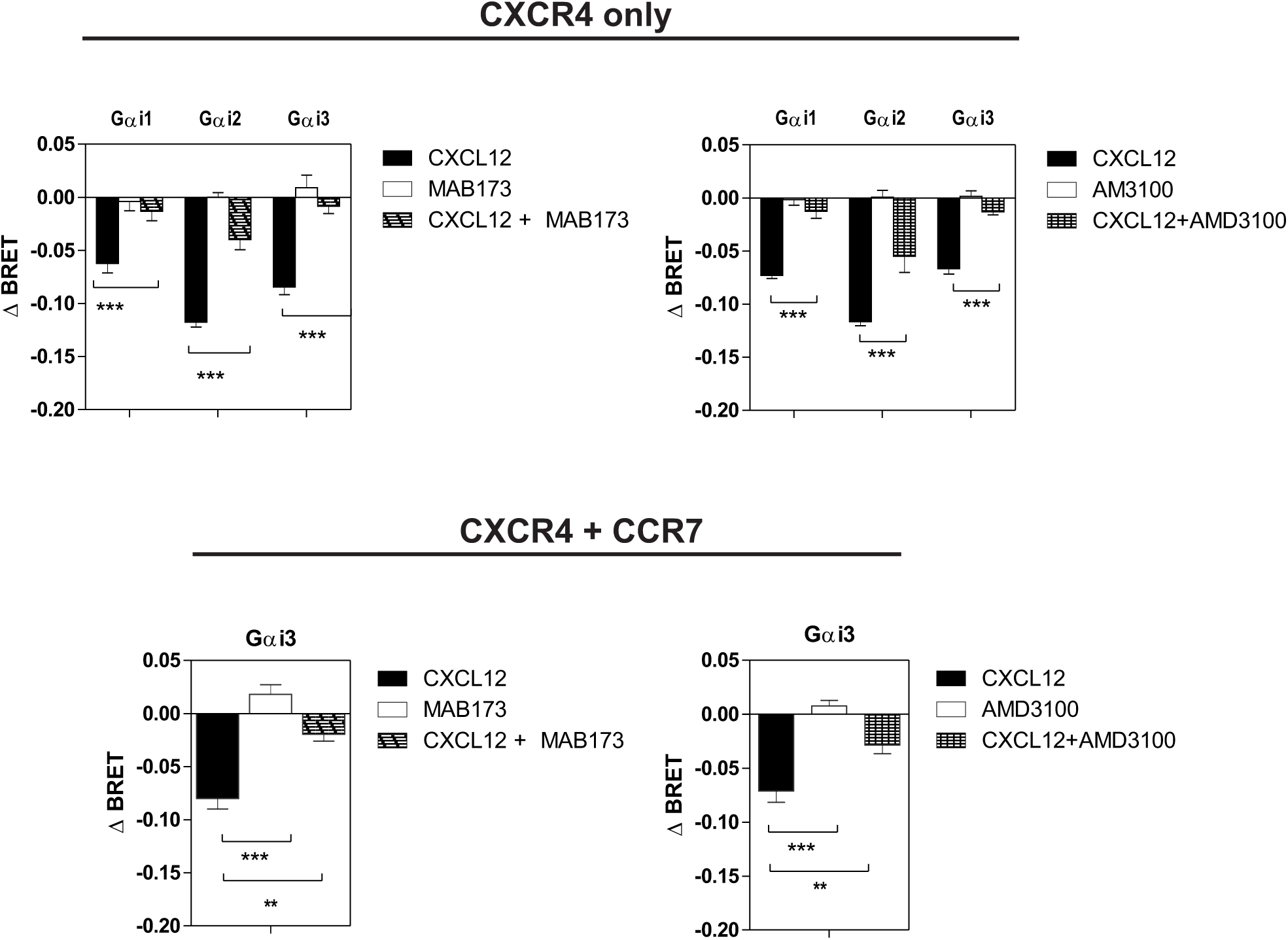
Activation of Gα proteins by CXCL12 is inhibited by CXCR4 antagonists. Real-time measurement of BRET signal in HEK293T cells coexpressing various Gαi biosensors with CXCR4 only (top) or in a combination with CCR7 (bottom). Cells were either left untreated or incubated 20 minutes with the anti-CXCR4 antibody MAB173 or the small molecule antagonist AMD3100 prior stimulation with 25 nM CXCL12. Results are expressed as ΔBRET corresponding to the difference in BRET signal between Gαi-*h*RLuc8 and Gβ1γ2-GFP10 measured in the presence and absence of CXCL12. Data are presented as mean values ± SEM (n=6; ** P<0.005, *** P<0,0005).

